# A central role of sibling sRNAs NgncR_162/163 in main metabolic pathways of *Neisseria gonorrhoeae*

**DOI:** 10.1101/2022.05.24.492095

**Authors:** Thomas Steiner, Marie Zachary, Susanne Bauer, Martin J. Müller, Markus Krischke, Sandra Radziej, Maximilian Klepsch, Bruno Huettel, Wolfgang Eisenreich, Thomas Rudel, Dagmar Beier

## Abstract

Bacterial regulatory RNAs (sRNAs) have been implicated in the regulation of numerous metabolic pathways. In most of these studies, sRNA-dependent regulation of mRNAs or proteins of enzymes in metabolic pathways has been predicted to affect the metabolism of these bacteria. However, only in very few cases has the role in metabolism been demonstrated. We performed here a combined transcriptome and metabolome analysis to define the regulon of the sibling sRNAs NgncR_162 and NgncR_163 and their impact on the metabolism of the major human pathogen *Neisseria gonorrhoeae*. These sRNA have previously been shown to control genes of the citric acid and methylcitrate cycle by post-transcriptional negative regulation. By transcriptome analysis we expand the NgncR_162/163 regulon by several new members and provide evidence that the sibling sRNAs act as both negative and positive regulators of target gene expression. Newly identified NgncR_162/163 targets are mostly involved in transport processes, especially the uptake of glycine, branched chain amino acids and phenylalanine. NgncR_162/163 also play key roles in the control of serine-glycine metabolism and hence probably affect biosynthesis of nucleotides, vitamins and other amino acids via the supply of C1-units. Metabolic flux analysis demonstrated a bipartite metabolism with glucose degradation providing intermediates for anabolic pathways, while energy metabolism via the citric acid cycle is mainly driven by amino acids, which feed into the cycle. Thus, by combined RNA-seq and metabolomics we significantly extended the regulon of NgncR_162/163 and demonstrate their role in the regulation of central metabolic pathways of the gonococcus.

**Importance:** *Neisseria gonorrhoeae* is a major human pathogen which infects more than 100 million people every year. An alarming development is the emergence of gonococcal strains resistant against virtually all of the antibiotics used for their treatment. Despite the medical importance and the vanishing treatment options of gonococcal infections, the bacterial metabolism and its regulation is only ill defined until today. We investigate here the regulation of the gonococcal metabolism by two previously studied sRNAs, NgncR_162/163 using RNA-seq and metabolomics. The results provided in this study demonstrate the regulation of transport processes and metabolic pathways involved in the biosynthesis of nucleotides, vitamins and amino acids by NgncR_162/163. Combined transcriptome and metabolome analyses provide a thus far unreached depth in the regulation of metabolic pathways by the neisserial sibling sRNAs and may therefore also be suitable for functional analysis of a growing number of other bacterial metabolic sRNA regulators.

## Introduction

*Neisseria gonorrhoeae* is the causative agent of the disease gonorrhoea, which is one of most common sexually transmitted infections. The World Health Organisation estimates 106 million new cases each year (WHO 2012). However, alarmingly not only the numbers of infections are rising, but also multidrug-resistant gonococcal strains are increasingly emerging, raising the threat of untreatable gonorrhoea (Rice et al., 2017). Gonococci infect mostly the mucosa of the urogenital tract, causing urethritis in men and cervicitis in women, but also pharynx, rectum and conjunctiva (Unema & Shafer, 2014). In rare cases, they can enter the bloodstream causing systemic disseminated gonococcal disease with manifestations like endocarditis, arthritis, dermatitis and sepsis (Rice et al., 2017).

The ability to infect different tissues requires effective regulatory mechanisms to ensure the pathogens rapid adaptation to changing environments. Non-coding RNAs have been recognized as an important class of post-transcriptional regulators of bacterial metabolism and virulence (Bobrovskyy et al., 2015; Gripenland et al., 2010). Trans-acting small RNAs (sRNAs) are typically transcribed from intergenic regions and undergo short imperfect base-pairing interactions with their target mRNAs. Most frequently, sRNA binding to the 5’-UTR of the target mRNA affects translation initiation by either obstructing the ribosome binding site (RBS) or by unfolding an intrinsic inhibitory secondary structure of the mRNA (Storz et al., 2011; Quereda et al., 2014). More recently, it was shown that negative regulation at the 5-UTR affecting translation initiation can also be accomplished by sRNA binding to ribosome standby sites or translational enhancer elements (Azam & Vanderpool, 2020; Yang et al., 2014; Darfeuille et al., 2007). Inhibition of translation is usually accompanied by increased ribonucleolytic target mRNA degradation in the absence of translating ribosomes. However, sRNA binding to both the 5′-UTR and the coding region can also directly impact mRNA half-live by either prevention of mRNA decay or stimulation of mRNA turnover by the active recruitment of ribonucleases (Storz et al., 2011; Papenfort & Vanderpool, 2015). Yet another mode of action of sRNAs is inhibition of Rho-dependent termination by blocking rut-sites, which were found to be frequently located in unusually long 5’-UTRs (Sedlyarova et al., 2016). Most studied sRNAs from Gram-negative bacteria require the interaction with an RNA chaperone like Hfq or ProQ in order to exert their function (Vogel & Luisi, 2011; Holmqvist et al., 2020).

Interestingly, multicopy sRNAs exhibiting a high degree of sequence identity, termed „sibling sRNAs”, were identified in some bacteria. Given their similarity, these sibling sRNAs may share a common seed region and therefore would be expected to act redundantly on the same target mRNAs. However, they may yet exert unique function due to differential expression of sRNA-encoding genes in response to particular environmental stimuli, by targeting different mRNAs via sequence motifs which are not conserved between the siblings or by mechanisms of gene regulation being unique to each sibling (reviewed in Caswell et al., 2014). The largest group of multicopy sRNAs known so far is represented by the LhrC family of *Listeria monocytogenes*, which comprises the highly homologous sRNAs LhrC1-5 as well as Rli22 and Rli33-1 exhibiting a lower degree of sequence conservation (Sievers et al., 2014; Mollerup et al., 2016). LhrC family sRNAs have an additive effect in target regulation, while the regulatory effect of a single sRNA seems to be rather subtle (Mollerup et al., 2016). Regulation of unique targets by sibling sRNAs is examplified by the AbcR sRNAs of *Sinorhizobium meliloti* (Torres-Quesada et al., 2013; Torres-Quesada et al., 2014).

A pair of sibling sRNAs has also been described in *N. gonorrhoeae* and *N. meningitidis* (Bauer et al., 2017; Heidrich et al., 2017; Pannekoek et al., 2017). These sRNAs, termed NgncR_162 and NgncR_163 in *N. gonorrhoeae*, are encoded adjacent to each other, exhibit 78% of sequence identity and fold into a similar secondary structure comprising three steem-loops. They are strongly expressed under standard laboratory growth conditions, however, NgncR_163 is considerably more abundant than NgncR_162. (Bauer et al., 2017). The *Neisseria* sibling sRNAs were shown to regulate the expression of genes involved in transcription regulation, amino acid uptake and basic metabolic processes including the methylcitrate and citric acid (TCA) cycle. All previously identified target genes are negatively regulated via hybridization of the sRNAs to the RBS of the target mRNA (Bauer et al., 2017; Heidrich et al., 2017; Pannekoek et al, 2017). Complete functional redundancy of the sibling sRNAs has been demonstrated experimentally (Bauer et al., 2017; Heidrich et al., 2017; Pannekoek et al., 2017) for a subset of targets.

To analyze the role of the sibling sRNAs in the neisserial metabolism in further detail, we combined transcriptome analysis and the investigation of carbon fluxes based on metabolomics and stable isotope incorporation experiments. By these approaches, several new target genes, including positively regulated targets, were identified and the sibling sRNAs were shown to interfere with sugar catabolism, the TCA cycle, serine-glycine metabolism and the transport of glycine, branched-chain amino acids and phenylalanine.

## Results

### Global transcriptome analysis reveals new members of the NgncR_162/163 regulon

We and others had identified target genes of the sibling sRNAs NgncR_162/163 and their homologues in *N. meningitidis* by *in silico* analysis of sRNA-mRNA interaction (Bauer et al., 2017; Heidrich et al., 2017) and comparative mass spectrometric analysis of cell lysates from wild-type bacteria and a sRNA double deletion mutant (Pannekoek et al., 2017). To obtain more complete insight into the NgncR_162/163 regulon, we compared the gene expression profile of *N. gonorrhoeae* wild-type MS11 and the sRNA double deletion mutant ΔΔ162/163 (Bauer et al., 2017) by deep RNA sequencing (RNA-seq). Genes differentially expressed in the two strains were assessed using DEseq2 (Love et al., 2014). A number of 95 protein coding genes (55 upregulated, 40 downregulated) exhibited expression ratios below 0.75 and above 1.5 (adjusted p-value (q) < 0.05) which was considered as differentially expressed (Table 1). Differentially expressed genes mostly belong to the functional categories (COG) “amino acid transport and metabolism” (14), “carbohydrate transport and metabolism” (7), “energy production and conversion” (7), “mobilome” (9) and “unknown function” (30). Putatively regulated ORFs encoding hypothetical proteins of unknown function tend to be very short and are most frequently located in the *N. gonorrhoeae* MS11 homologues of the *maf* genomic islands of *N. meningitidis* encoding secreted polymorphic toxins (*mafB*), their specific immunitiy proteins (*mafI*) as well as alternative MafB C-terminal domains (MafB-CT) or MafI modules (Jamet et al., 2005). From the NgncR162/163 target genes validated previously by qRT-PCR (Bauer et al., 2017) NGFG_01721, NGFG_02049, *prpC, ack* and *gltA* were detected by RNA-seq, whereas *prpB, gdhR, fumC* and *sucC* did not meet the applied cut-off, reflecting differences in the sensitivity of the two RNA quantification techniques.

**Table 1:**
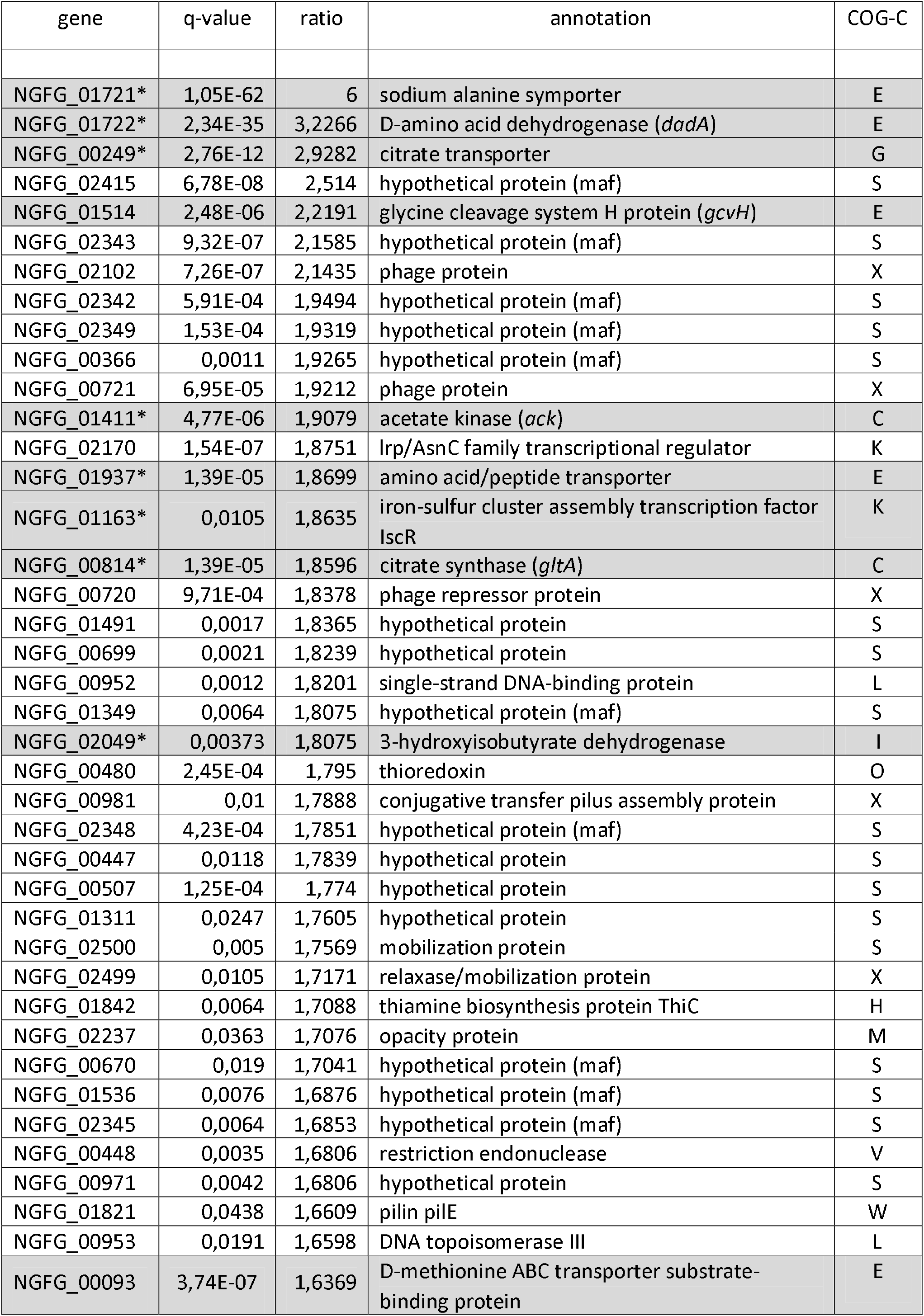
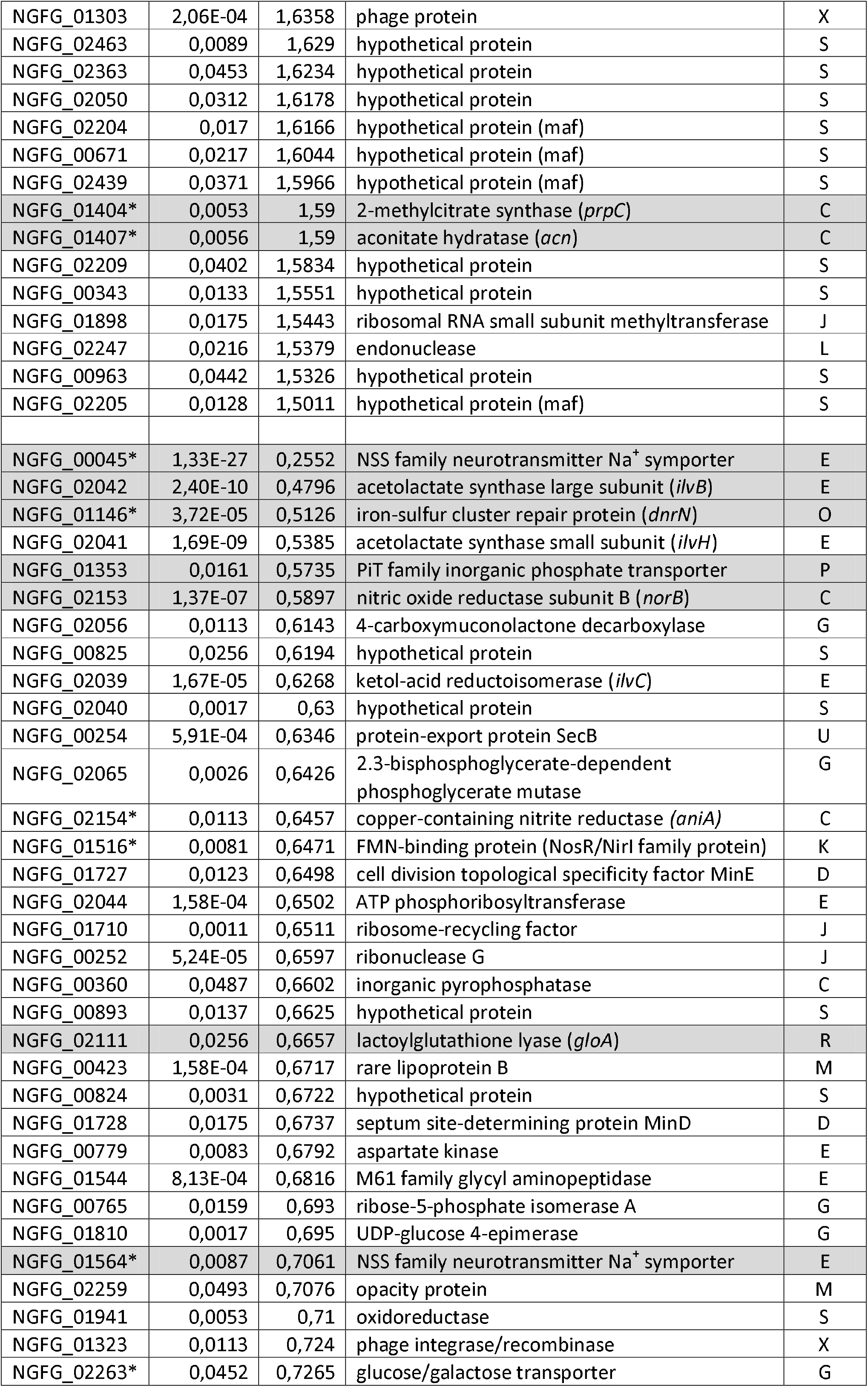
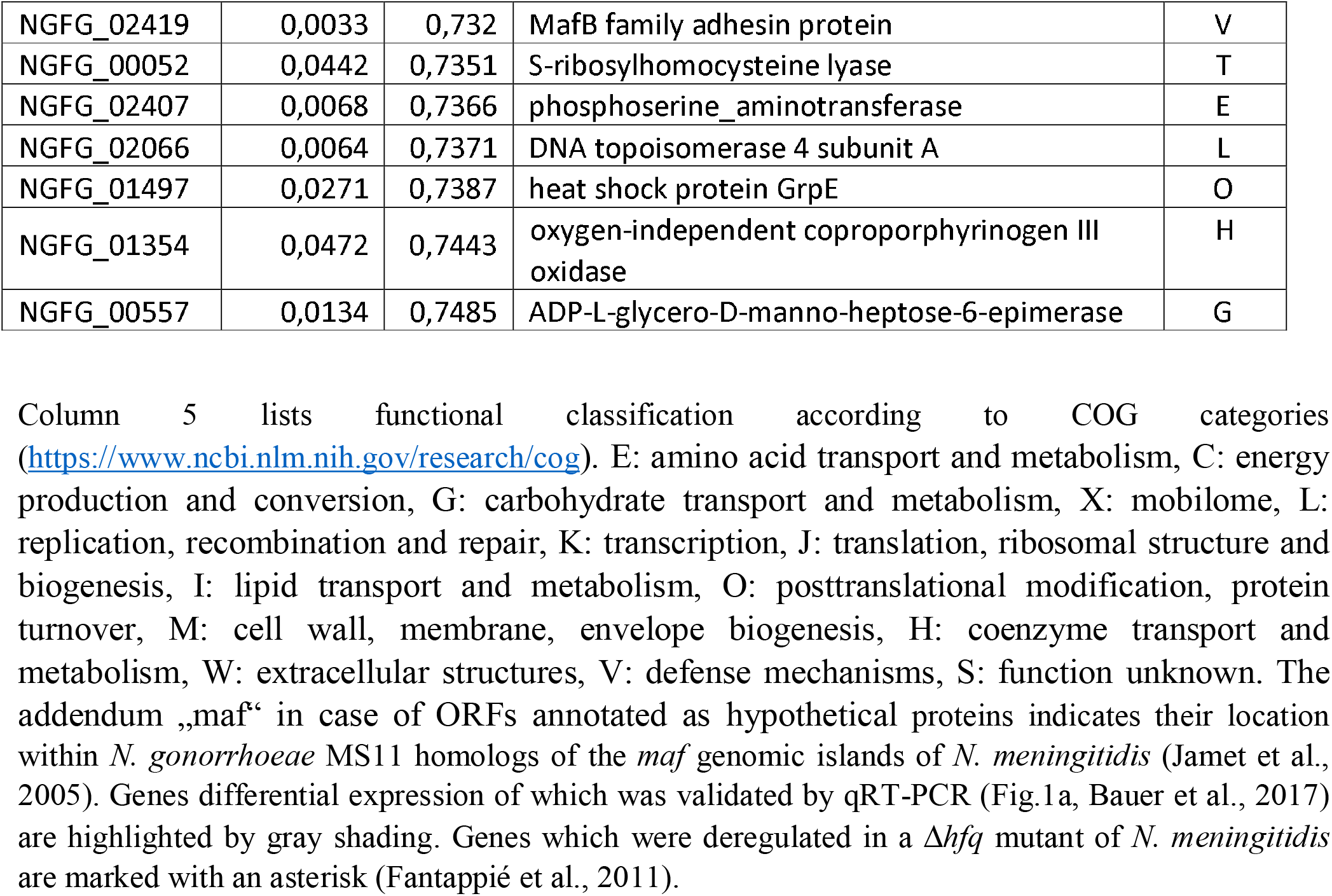
Genes differentially expressed in MS11 ΔΔ162/163 within a <0,75 and >1.5 cut-off for fold expression change compared to MS11 (q <0.05). RNA-seq was performed with samples from three biological replicates each.

15 putative NgncR_162/ 163 target genes with predicted metabolic function were selected for target validation via qRT-PCR analysis (Fig. 1a) in mutant ΔΔ162/163 and the complemented mutant (ΔΔc162/163) carrying both sRNAs genes integrated in the *iga*-*trpB* locus (Bauer et al., 2017). With exception of sugar transporter NGFG_02263 differential expression in mutant ΔΔ162/163 could be confirmed. NGFG_02042 encoding *ilvB* from the branched chain amino acid (BCAA) biosynthetic pathway was significantly downregulated in ΔΔ162/163, however expression was not reverted in the complemented mutant ΔΔc162/163. Validated target genes encode proteins involved in the transport of amino acids or peptides (NGFG_00045, NGFG_01564, NGFG_00039, NGFG_01937), citrate (NGFG_00249) or inorganic phosphate (NGFG_01353), as well as NGFG_01722 being cotranscribed with the putative alanine symporter NGFG_01721 and encoding *D*-alanine dehydrogenase (*dadA*). Furthermore, differential expression of lactate permease (NGFG_01471) with a ratio slightly above the applied cut-off in RNA-seq (0.765779; q = 0,0191) could be validated by qRT-PCR (Fig. 1a). Another validated target, aconitate hydratase (*acn*) belongs to the methylcitrate cycle, which was previously shown to be controlled by the sibling sRNAs (Bauer et al., 2017; Heidrich et al., 2017). Interestingly, NGFG_01514 encoding *gcvH* was found to be strongly upregulated in the absence of the sibling sRNAs. GcvH is a component of the glycine cleavage system which converts glycine to CO_2_, NH_3_ and 5,10-methylene-tetrahydrofolate (5,10-MTHF), suggesting a role of the sibling sRNAs in purine, histidine, thymine, panthothenate and methionine synthesis via control of the supply of C1-units. Furthermore, RNA-seq suggests a contribution of the sibling sRNAs in the control of iron-sulfur cluster synthesis and homeostasis (*iscR, dnrN*) and anaerobic respiration (*aniA, norB*, NGFG_01516). Notably, 16 from the putative targets of the sibling sRNAs listed in Table 1 as well as lactate permease were also differentially expressed in a *hfq* deletion mutant of *N. meningitidis* (Fantappié et al., 2011). Since the sibling sRNAs were shown to bind Hfq in both *N. meningitidi*s and *N. gonorrhoeae* and their stability was largely diminished in its absence (Heidrich et al., 2017; Heinrichs and Rudel, unpublished), deregulation of transcript levels in a *hfq* mutant is likely to be the consequence of post-transcriptional regulation mediated by the sibling sRNAs. In case of NGFG_01471, differential expression in ΔΔ162/163 can be explained by an indirect effect, because lactate permease was shown to be regulated by GdhR (Ayala & Shafer, 2019), which itself is a target of the sibling sRNAs (Bauer et al., 2017).

**Fig. 1.**
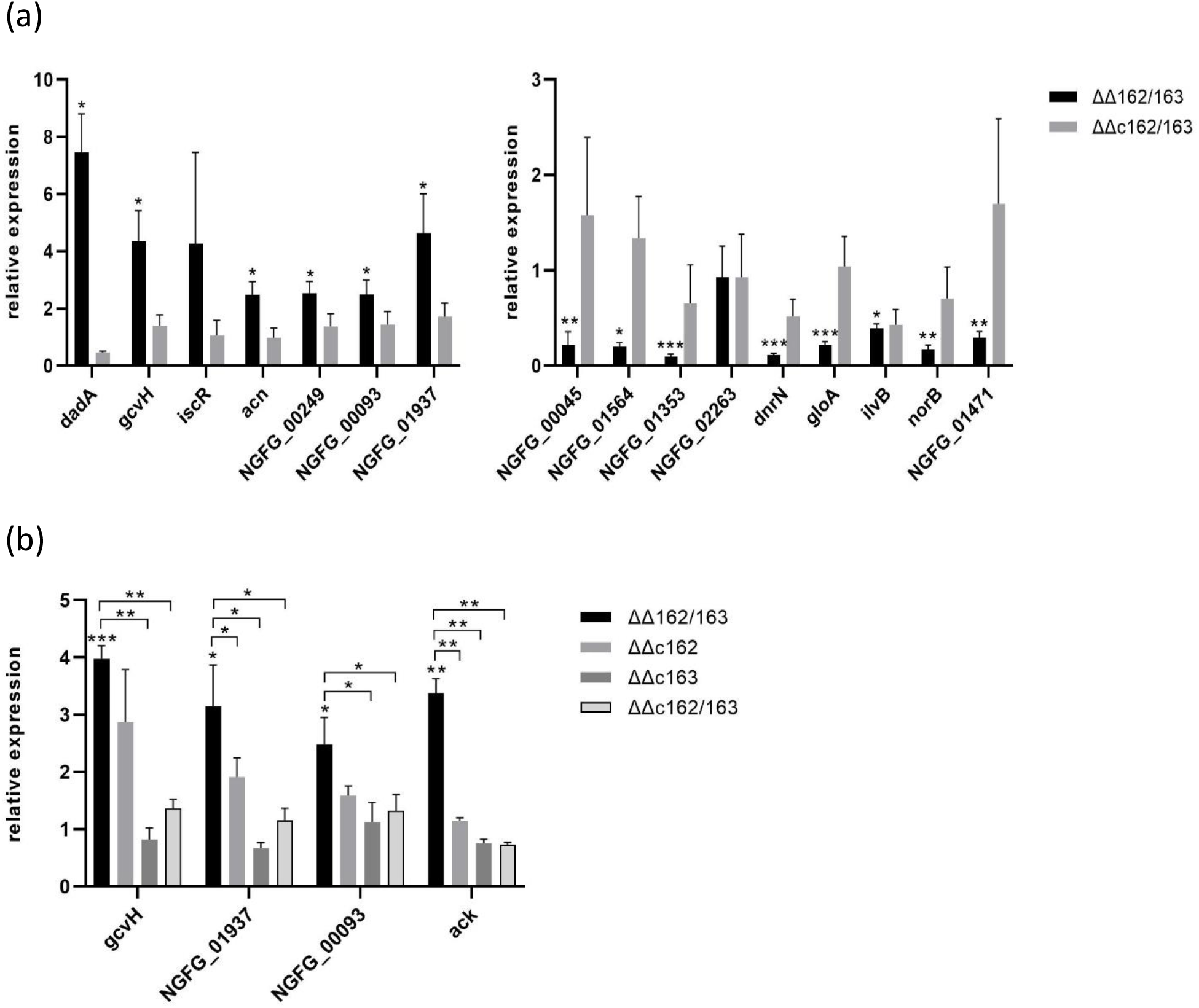
(a) Validation of putative target genes, which were found differentially expressed in the absence of the sibling sRNAs in RNA-seq analysis. Transcript levels of target genes were analysed by qRT-PCR in strains MS11, the sRNA double deletion mutant ΔΔ162/163 and the complemented strain ΔΔc162/163. The ratios of the transcript amount relative to the wild-type MS11 (normalized to 1) are depicted. (b) In case of *gcvH*, NGFG_00093 and NGFG_01937 transcript levels were also assessed in derivatives of MS11 ΔΔ162/163 complemented with either NgncR_162 (ΔΔc162) or NgncR_163 (ΔΔc163). *ack* was used as a control since the sibling sRNAs were previously shown to display functional redundancy on this target (Bauer et al., 2017). The indicated ratios represent the mean of the results of qRT-PCR experiments performed in triplicate on cDNAs obtained from at least three independent RNA preparations. Error bars indicate the standard deviation. Statistical significance was determined using Student’s t-test analysis (*=p<0.05, **=p<0.01, ***=p<0.001).

To test functional redundancy of NgncR_162 and NgncR_163 in the regulation of newly identified targets, we exemplarily investigated the impact of the individual sibling sRNAs as post-transcriptional regulators of *gcvH*, NGFG_01937 and NGFG_00093. Complementation of ΔΔ162/163 with NgncR_163 restored mRNA abundance to wild-type levels, while only partial complementation was observed in case of NgncR_162 (Fig. 1b). These data suggest that the individual siblings act in a hierarchical manner on certain targets, while exhibiting complete functional redundancy on others.

### sRNA-target interaction is predicted to occur within the 5’-UTR and coding region of the regulated mRNAs

All targets identified previously (Bauer et al., 2017; Pannekoek et al., 2017, Heidrich et al., 2017) are negatively regulated by the sibling sRNAs via mRNA-sRNA duplex formation resulting in obstruction of the RBS. sRNA-mRNA interaction involved the single-stranded region of stem-loop (SL) 2 and in some cases the single-stranded region connecting SL1 and SL2 (SSR1). Complementarity between the validated new target mRNAs and the sibling sRNAs was analysed using IntaRNA (Mann et al., 2017) (Fig. S1). In accordance with negative regulation of *dadA*, NGFG_00249, NGFG_01937 and NGFG_00093 base pairing interactions involving the SL2 loop or SSR1 sequence, which mask the RBS, were predicted for these targets. Surprisingly, also in case of the positively regulated targets *dnrN* and *norB* a region of complementarity to SL2 overlapping the RBS was detected. Other members of the NgncR_162/163 regulon being under negative or positive control are targeted within the coding sequence (CDS). In case of NGFG_01564 and *gcvH* mRNAs, short (8-9 nt) segments with base-pairing capability to the sRNA SSR1 or SL2 sequence are present, while other targets exhibit more extended regions of partial complementarity, which engage both SSR1 and SL2 sequence (*acn, iscR*, NGFG_00045) or both SL2, the single-stranded region connecting SL2 and SL3 (SSR2) and part of the SL3 sequence (*gloA*) of both NgncR_162 and NgncR_163. Both the SL1 and SSR1 sequence of NgncR_162 show partial complementarity to a 28-nucleotides region within the CDS of NGFG_01353. Due to sequence variation in SL1 a similar interaction is not predicted for NgncR_163. However, a second interaction site is predicted in NGFG_01353 with complementarity to the SL2 and SSR2 sequence of both sRNAs. Thus, putative hybridization regions can be predicted for most of the identified target mRNAs.

### NGFG_00045 mRNA level is directly affected by the sibling sRNAs

RNAs downregulated in the absence of NgncR_162/163 identified in this study represent a new class of targets. To demonstrate the validity of this class of targets, transporter NGFG_00045 was chosen to investigate positive regulation by the sibling sRNAs in more detail. Northern blot analysis was performed on RNA extracted from MS11, ΔΔ162/163 and the complemented mutant ΔΔc162/163. RNA-seq had revealed that NGFG_00045 is cotranscribed with a gene encoding a hypothetical peptide of 30 amino acids (Remmele et al., 2015) and a transcript of the expected size was observed in wild-type MS11 and the complemented mutant, but was barely detectable in ΔΔ162/163 (Fig. 2a). The hybridization pattern of RNA from mutant ΔΔ162/163 was unaltered, indicating that the sibling sRNAs are not involved in prominent processing events (Fig. 2a). Next, we replaced the promoter region of NGFG_00045 (Remmele et al., 2015) with the promoter of a gonococcal *opa* gene in MS11, ΔΔ162/163 and ΔΔc162/163. qRT-PCR and Northern blot analysis showed that transcript amounts were still affected by the absence of the sibling sRNAs (Fig. 2b, c). When on the contrary promoter region and 5’-UTR of NGFG_00045 were fused to *gfp*, almost equal amounts of mRNA were detected in the presence or absence of the sibling sRNAs (Fig. 2d). Since expression of GFP protein could not be detected in these mutants due to a weak RBS in the NGFG_00045 5’-UTR, we introduced the consensus *E. coli* Shine-Dalgarno sequence via site directed mutagenesis. As observed for *gfp* mRNA, protein levels were not affected in the absence of the sibling sRNAs (Fig. 2e). These data demonstrated that positive regulation of NGFG_00045 is not indirectly mediated via a transcriptional regulator being the target of the sRNAs and that the 5’-UTR of NGFG_00045 is not involved in post-transcriptional regulation. Attempts to investigate whether mRNA stability is affected by the absence of the sibling sRNAs by Northern blot analysis of RNA extracted from rifampicin-treated cultures were hampered by the fact that NGFG_00045 mRNA amounts are extremely low in strain ΔΔ162/163. The mechanism of NGFG_00045 target activation by the sibling sRNAs remains to be investigated in more detail.

**Fig. 2.**
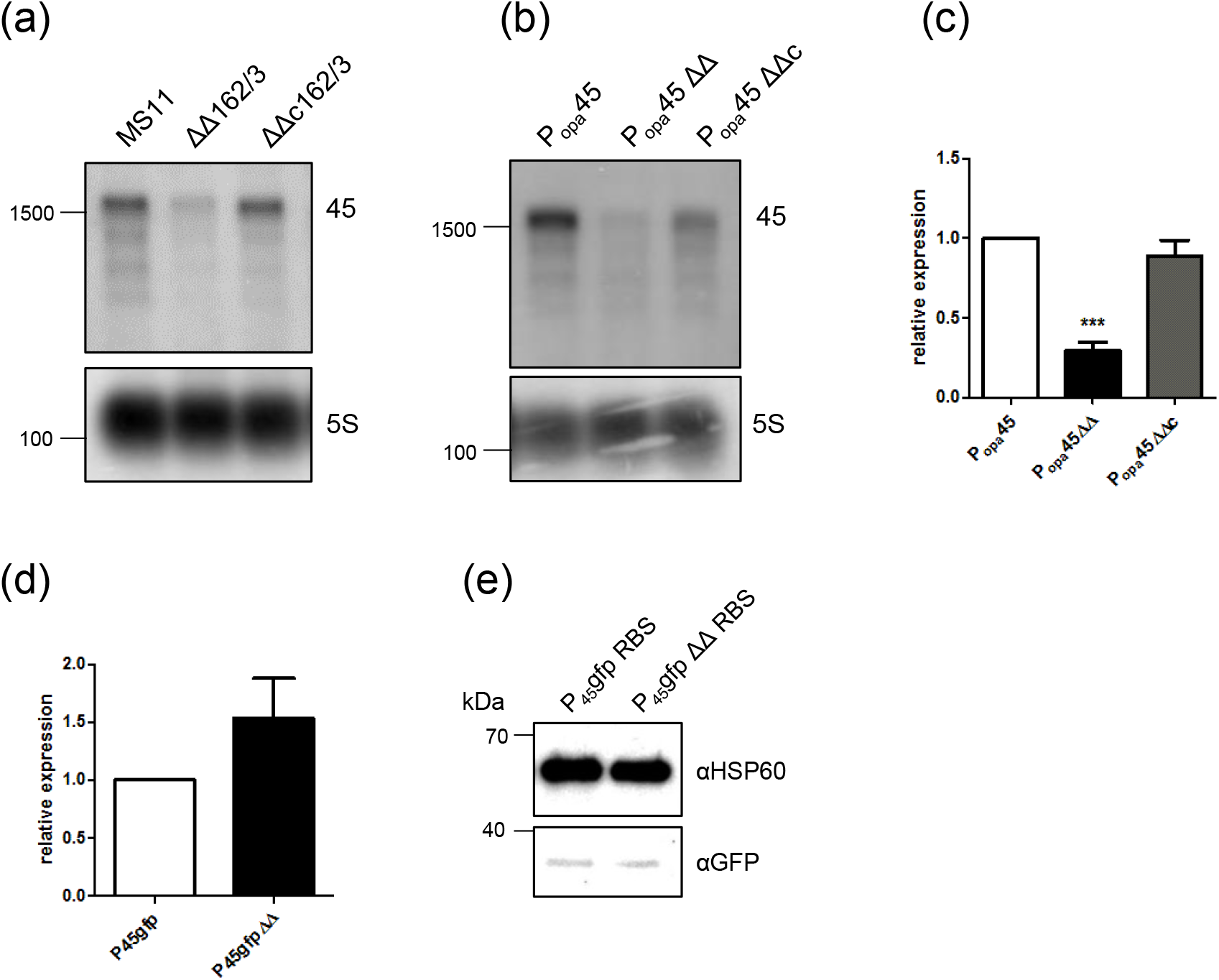
Expression analysis of NGFG_00045 in the presence and absence of the sibling sRNAs. (a) and (b) Equal amounts of RNA extracted from MS11, ΔΔ162/163 and ΔΔc162/163 (a) or P_*opa*_45, P_*opa*_45ΔΔ and P_*opa*_45ΔΔc (b) grown to logarithmic phase were analysed by Northern blotting using a NGFG_00045-specific radiolabelled probe which was generated by PCR using primer pair qRT45-1/qRT45-2 (Table S2). Probing for 5S rRNA was used as loading control. The position of radiolabelled marker RNAs of 1500 and 100 nucleotides is indicated on the left side of the panels. (c) Transcription of NGFG_00045 under control of the P_*opa*_ promoter in strains P_*opa*_45, P_*opa*_45ΔΔ and P_*opa*_45ΔΔc was analysed by qRT-PCR. The ratios of the transcript amount relative to P_*opa*_45 (normalized to 1) are depicted. The indicated ratios represent the mean of the results of qRT-PCR experiments performed in triplicate on cDNAs obtained from three independent RNA preparations. Error bars indicate the standard deviation. Statistical significance was determined using Student’s t-test analysis (***=p<0.001). (d) Transcription of the reporter gene *gfp* in strains P_45_gfp and P_45_gfpΔΔ expressing *gfp* under control of the NGFG_00045 promoter was quantified by qRT-PCR. The transcript amount in P_45_gfp was normalized to 1. The indicated ratio represents the mean of the results of qRT-PCR experiments performed in triplicate on cDNAs obtained from two independent RNA preparations. (e) Western blot analysis was performed on equal amounts of protein extracted from strains P_45_gfpRBS and P_45_gfpΔΔRBS with monoclonal antibodies directed against GFP and HSP60 used as loading control. Numbers on the left side of the panel indicate the position of size marker proteins.

### Amino acid uptake is altered in ΔΔ162/163 and a NGFG_00045 deletion mutant

Since several targets of the sibling sRNAs were predicted to be involved in amino acid transport, we first analysed amino acid uptake in wild-type, ΔΔ162/163 and complemented mutant to identify changes associated with sibling sRNA expression. Bacteria were grown in chemically defined medium to logarithmic phase and amino acid levels in the culture supernatants were analysed by mass spectrometry. Growth of wild-type MS11 resulted in 90% depletion of glutamine and glutamate, and about 50% and 40% reduction of proline and asparagine concentrations, respectively, in the spent culture medium (Fig. S2). Consumption was marginal in case of glycine, alanine, arginine, tyrosine, tryptophan, valine and histidine (less than 5% of the supplied amounts) and moderate (ranging from 5 to 30%) for other amino acids. Surprisingly, the concentration of aspartate seemed to be even increased in the culture supernatant (Fig. S2), suggesting lack of aspartate transport, which might be compensated by efficient biosynthesis of oxaloacetate from phosphoenolpyruvate (PEP) via PEP carboxylase (Cox et al., 1977; see below). Compared to the wild-type, the sRNA double-mutant ΔΔ162/163 showed significant differences in the consumption of proline, glycine, alanine, serine and threonine. More specifically, glycine, alanine and proline consumption was increased 9-, 4- and 1.2-fold, respectively, while serine and threonine uptake was diminished about 1.5-fold. As expected, the amino acid profile of the culture supernatant of the complemented strain ΔΔc162/163 resembled that of the wild-type (Fig. 3).

**Fig. 3.**
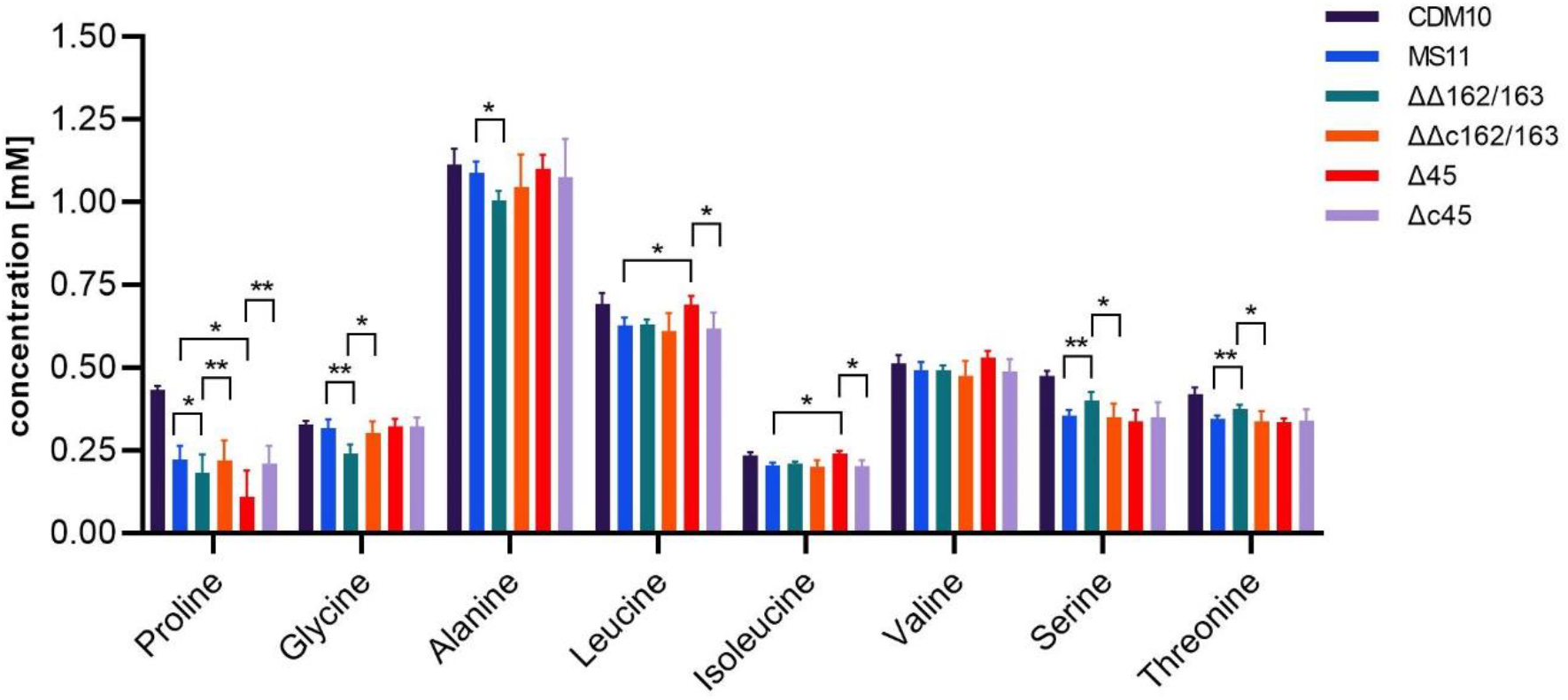
Analysis of the impact of sRNA and NGFG_00045 knockout on amino acid uptake by *N. gonorrhoeae. N. gonorrhoeae* MS11, ΔΔ162/163, ΔΔc162/163, Δ45 and Δc45 were grown to OD_550_ = 0.6 in modified CDM10 medium. Bars depict the amino acid concentration (mM) of culture supernatants of *N. gonorrhoeae* strains determined by mass spectrometry and represent the mean of four biological replicates (5 technical replicates each). For comparison amino acid concentrations determined for CDM10 medium (see Fig. S2) are included. Error bars indicate the standard deviation. Statistical significance was determined using Student’s t-test analysis (*=p<0.05, **=p<0.01).

To investigate the contribution of sibling sRNA targets to amino acid transport, knockout mutants for NGFG_01721, NGFG_00045 and NGFG_01564 were created. In the NGFG_00045 knockout mutant Δ45, proline uptake was massively increased, while the uptake of leucine, isoleucine and valine was abolished. Again, the observed effects were reversed in the respective complementation mutant Δc45 (Fig. 3). Analysis of the amino acid composition of culture supernatants of mutants Δ1721 and Δ1564 did not reveal statistically significant differences to wild-type MS11(data not shown). In Δ45, increased consumption of proline, which can be efficiently converted to glutamate via the bifunctional proline dehydrogenase/pyrroline-5-carboxylate dehydrogenase NGFG_01376, apparantly compensates for the amino acid uptake defect caused by the inactivation of NGFG_00045, which is likely to be a branched-chain amino acid (BCAA) transporter. Elevated proline uptake in ΔΔ162/163 is consistent with downregulation of NGFG_00045 in this mutant. Furthermore, increased glycine uptake in ΔΔ162/163 strongly argues in favour of NGFG_01721 being predominantly a glycine transporter, since this protein is massively upregulated in the absence of the sibling sRNAs (Table 1; Bauer et al., 2017). Glycine cleavage, which is affected by the sibling sRNAs via target *gcvH*, yields 5,10-MTHF, which together with another molecule of glycine can then be converted to serine by the serine hydroxymethyltransferase GlyA (reviewed in Stauffer, 2004) (Fig. 7). In *N. meningitidis, glyA* expression was reported to be derepressed in the absence of *hfq* and the sibling sRNAs, respectively (Fantappié et al., 2011; Pannekoek et al., 2017). Since serine uptake was diminished in mutant ΔΔ162/163 and in our RNA-seq analysis *glyA* was weakly upregulated with a q-value close to significance (fold change:1.29; q = 0.059), we validated *glyA* transcript levels by qRT-PCR. In fact, upregulation of *glyA* was observed in the sRNA double deletion mutant, while wild-type transcript level was restored in the complemented strain (Fig. S3a). Furthermore, a translational *glyA*-*gfp* fusion was downregulated in the presence of NgncR_162 in *E. coli* (Fig. S3b), confirming negative regulation of *glyA* by the sibling sRNAs. This is in accordance with predicted binding of the sibling sRNAs to the RBS of the *glyA* mRNA (Fig. S1). Therefore, we conclude that derepression of NGFG_01721, *gcvH* and *glyA* in the sRNA double-mutant enhances serine biosynthesis resulting in a decreased demand for serine uptake.

### Isotopologue profiling reveals a bipartite metabolism in *N. gonorrhoeae*

From their target spectrum as well as measurements of amino acid concenctrations in the culture supernatant, it became apparent that the sibling sRNAs in *N. gonorrhoeae* are involved in the regulation of the central carbon metabolism in response to available nutrients in the environment. The central carbon metabolism in *N. gonorrhoeae* has so far only been investigated using enzyme assays with cell extracts or genome annotation (Morse et al., 1974; Chung et al., 2008). To better define the central carbon metabolism of *N. gonorrhoeae* and investigate the actual metabolic phenotype during growth as well as the effects of the sibling sRNAs, we herein employed stable isotope incorporation experiments with subsequent isotopologue profiling.

To this aim, the bacteria were grown for 4 h in duplicate to logarithmic phase (OD_550_ = 0.6) in chemically defined medium containing fully ^13^C-labelled glucose or proline ([U-^13^C_6_]glucose or [U-^13^C_5_]proline). After harvest, the cells were mechanically disrupted (fraction 1) or hydrolysed under acidic conditions (fraction 2). Protein derived amino acids (in fraction 2) and fatty acids (in fraction 1) were silylated and then applied to GC-MS analysis (three technical replicates). From the relative masses detected for the specific fragments in the MS spectra, the ^13^C-excess values of amino acids and fatty acids were calculated using the software package Isotopo (Ahmed et al., 2014). Here, the overall ^13^C-excess values show ^13^C-contents beyond the natural ^13^C abundances in the respective molecules. Moreover, using the same software the isotopologue compositions were determined displaying the relative fractions (%) of isotopologues (M+1, M+2, M+3, …, M+n) for each molecule under study. Herein, M denotes the molecular mass with only ^12^C in the carbon backbone of the molecule and n specifies the number of ^13^C-carbons.

Using 13.9 mM [U-^13^C_6_]glucose as supplement to the medium, high ^13^C-excess values were detected in alanine (41.3% ^13^C-excess) with about 90% M+3 (i.e. displaying the fraction of the [U-^13^C_3_]- species) and valine (44.7% ^13^C-excess) with about 70% M+5 (Fig. 4a and b). This confirmed the efficient uptake and utilization of [U-^13^C_6_]glucose to afford fully labelled pyruvate, which was then converted into the detected [U-^13^C_3_]alanine and [U-^13^C_5_]valine specimens, respectively (see also Fig. 4c). Although the labelling patterns from [U-^13^C_6_]glucose do not allow a distinction between glycolysis and the Enter-Doudoroff (ED) pathway for glucose degradation, the ED pathway is the predominant pathway for pyruvate formation from glucose in *N. gonorrhoeae*, as based on enzyme assays (Morse et al., 1987). In addition, glycolysis seems to be non-functional due to the absence of the gene for phosphofructokinase in the genome of *N. gonorrhoeae* (Chung et al., 2008).

**Fig. 4.**
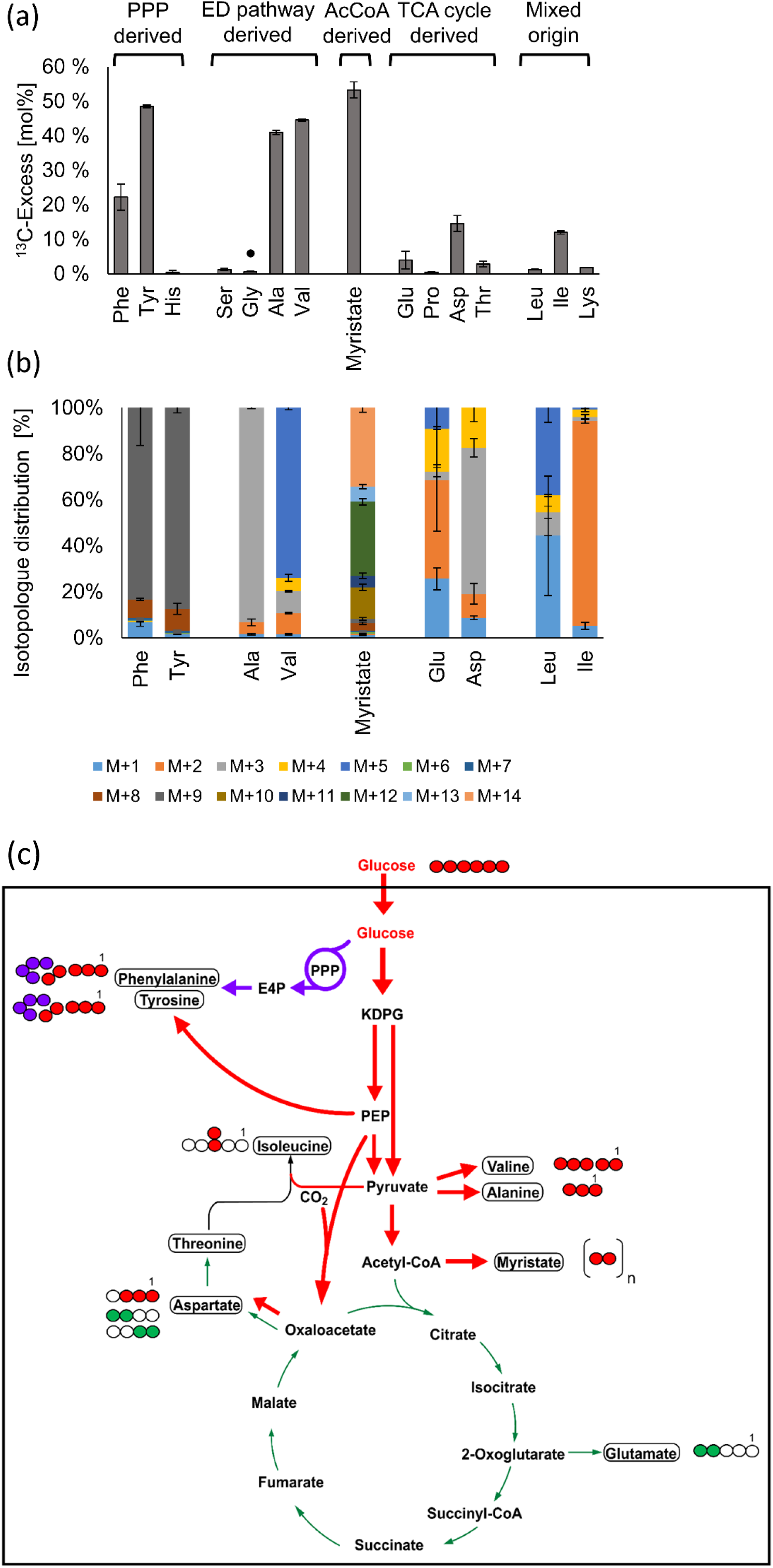
^13^C-Excess and isotopologue profiles of myristate and protein-derived amino acids from *N. gonorrhoeae* after growth in the presence of 13.9 mM [U-^13^C_6_]glucose. (a) The ^13^C-excess (mol%) displays the overall ^13^C-incorporation in the respective metabolite beyond natural ^13^C-abundance. The amino acids are arranged in groups according to their relation to the central carbon metabolism. The values are means from two biological replicates and three technical replicates. The error bars indicate mean deviations. Glycine is not derived from a C3-intermediate, but rather formed from threonine (for details, see text) (b) Isotopologue compositions of myristate and amino acids with siginficant ^13^C-excess; M denotes the molecular mass with only ^12^C in the carbon-backbone of the respective amino acid, while M+X represents the incorporation of X ^13^C-atoms. The values are means from two biological replicates and three technical replicates. The error bars indicate mean deviations. (c) Scheme of the carbon fluxes starting from the supplied [U-^13^C_6_]glucose in the central carbon metabolism of *N. gonorrhoeae*. The red arrows indicate fluxes into PEP and pyruvate via the ED pathway and into acetyl-CoA for fatty acid biosynthesis. Downstream reactions from these intermediates into amino acids and oxaloacetate are also indicated by red arrows. The purple arrows indicate fluxes via the PPP and the shikimate pathway, and the green arrows indicate fluxes from acetyl-CoA into the TCA and the downstream amino acids, glutamate, aspartate and threonine via citrate, 2-oxoglutarate and oxaloacetate, respectively. The thickness of the arrows corresponds to the approximate relative extent of ^13^C-incorporation from the labelled tracer. The boxes indicate metabolic products which were analysed by GC-MS. The filled circles indicate ^13^C-labelled positions in these molecules, whereas the open circles indicate unlabelled positions.

Tyrosine (^13^C-excess of 48.6%) and phenylalanine (^13^C-excess of 19.5%) also reflected the efficient incorporation of exogenous ^13^C-glucose via the ED pathway and the pentose phosphate pathway (PPP) producing M+4 erythrose-4-phosphate (E-4-P) and M+3 phosphoenolpyruvate (PEP) as precursors. More specifically, the formation of the detected M+9 isotopologues in tyrosine and phenylalanine can be easily explained by the assembly of the aromatic amino acids from [U-^13^C_4_]E-4-P and two molecules of [U-^13^C_3_]PEP via the shikimate pathway (Fig. 4c). These observations are in full accordance with the reported genome sequence (Chung et al., 2008) and enzyme assays of sugar degrading pathways (Morse et al., 1987).

Although serine is typically derived from 3-phosphoglycerate via 3-hydroxypyruvate, the genome of *N. gonorrhoeae* does not encode an enzyme for the 3-phosphoglycerate reduction into 3-hydroxypyruvate. Indeed, only a minor ^13^C excess (1.4%) was observed for serine from the protein hydrolysate in the [U-^13^C_6_]glucose experiment (Fig. 4). In conclusion, serine was mainly derived from unlabelled serine and other unlabelled components in the medium. This was also in accordance with high serine uptake observed through culture supernatant analysis (Fig. S2). Not surprisingly then, almost no ^13^C-excess was detected in glycine (0.7%), since glycine was derived from serine or was also directly taken up from the medium in unlabelled form. Similarly, due to the absence of histidinol phosphatase in the genome sequence, no ^13^C-excess was detected in histidine.

The high ^13^C-excess of the fatty acid myristate (about 55%) reflected the conversion of M+3 pyruvate into M+2 acetyl-CoA, a precursor of fatty acid biosynthesis. Incorporation of M+2 acetyl-CoA as a building block was detected as a mixture of ^13^C-isotopologues in myristate mainly with even numbers of ^13^C-atoms (i.e. M+8, M+10, M+12 and M+14) indicating fatty acid biosynthesis using 4, 5, 6 or 7 [U-^13^C_2_]acetyl-CoA units, respectively (Fig. 4b). A lower amount of M+2 labelled acetyl-CoA was channeled into the oxidative TCA cycle (Hebeler et al., 1976; Chung et al., 2008) via synthesis of citrate, which was then metabolized to [4,5-^13^C_2_]2-oxoglutarate and [4,5-^13^C_2_]glutamate (3.9%) (Fig. 4). In contrast, a higher ^13^C-enrichment occurred in aspartate (14.6%) mainly due to the high relative fractions of M+3 (about 60%) as well as M+4 (about 20%) (see Fig. 4). Based on these results, it can be assumed that PEP carboxylase (Cox et al., 1977) was highly active converting [U-^13^C_3_]PEP and ^12^CO_2_ or ^13^CO_2_ into [1,2,3-^13^C_3_]oxaloacetate or [U-^13^C_4_]oxaloacetate, which were further converted to the observed [1,2,3-^13^C_3_]- and [U-^13^C_4_]aspartate species, respectively (Fig. 4b). ^13^CO_2_ was potentially produced directly in the cell during decarboxylation of [U-^13^C_3_]pyruvate to [1,2-^13^C_2_]acetyl-CoA and could therefore locally have produced a ^13^C-enrichment in CO_2_ beyond natural abundance. Notably, only small amounts of aspartate seemed to be synthetized via the TCA cycle, since the TCA-derived ^13^C_2_-aspartate accounted for only 10% of its isotopologue profile (Fig 4b).

Additionally, some ^13^C-label was observed in isoleucine (12 %) by incorporation of a labelled pyruvate unit into C-3 and the attached methyl group (Fig 4). No significant ^13^C-excess was detected in leucine (1.1 %), threonine (2.2 %), lysine (1.8 %), and proline (0.5%), although enzymes for the biosynthesis of proline from glutamate were annotated in the genome (Chung et al., 2008).

Since only low ^13^C-enrichments were detected in amino acids using intermediates of the TCA cycle in these experiments, we assumed that the TCA in *N. gonorrohoeae* is mainly fuelled by unlabelled compounds from the medium (i.e. not from [U-^13^C_6_]glucose). The ΔΔ162/163 mutant had revealed an increased uptake of proline (Fig. 3), which indicated that this amino acid could serve as a precursor for glutamate and TCA cycle intermediates via 2-oxoglutarate. Therefore, a labelling experiment starting from [U-^13^C_5_]proline in the medium was performed. Indeed, [U-^13^C_5_]proline added to the culture medium was efficiently taken up and utilized leading to 80% M+5 in the isotopologue profile of glutamate (17.7 % ^13^C-excess) and about 80% M+4 in the isotopologue profiles of aspartate (20% ^13^C-excess) and threonine (5.7% ^13^C-excess) (Fig 5). In contrast, only low ^13^C-excess was detected in alanine, glycine, valine and isoleucine (below 5%), whereas serine did not contain ^13^C beyond the natural abundance content. Based on this observation, labelled glycine could hardly be formed form the apparently unlabelled serine. Rather, glycine was potentially synthesized from (labelled) threonine via the threonine utilisation (Tut) pathway (reviewed in Stauffer, 2004) as the genome of *N. gonorrhoeae* contains distant homologues of threonine dehydrogenase and 2-amino-3-ketobutyrate CoA ligase from *Escherichia coli*.

The different ^13^C-enrichments from the labelling experiments with ^13^C-glucose and ^13^C-proline indicated that *N. gonorrhoeae* used different substrates simultaneously for its growth suggesting a bipartite metabolic network as shown in Fig. 6. In this model, glucose is used as a substrate providing precursors for anabolic purposes, i.e. for sugar components and some aromatic amino acids derived from the PPP, amino acids derived from pyruvate and fatty acids derived from acetyl-CoA. In contrast, glucose does not serve as a major substrate to feed the TCA cycle. Rather, proline or related compounds drive the TCA cycle and mainly contribute to energy metabolism.

**Fig. 5.**
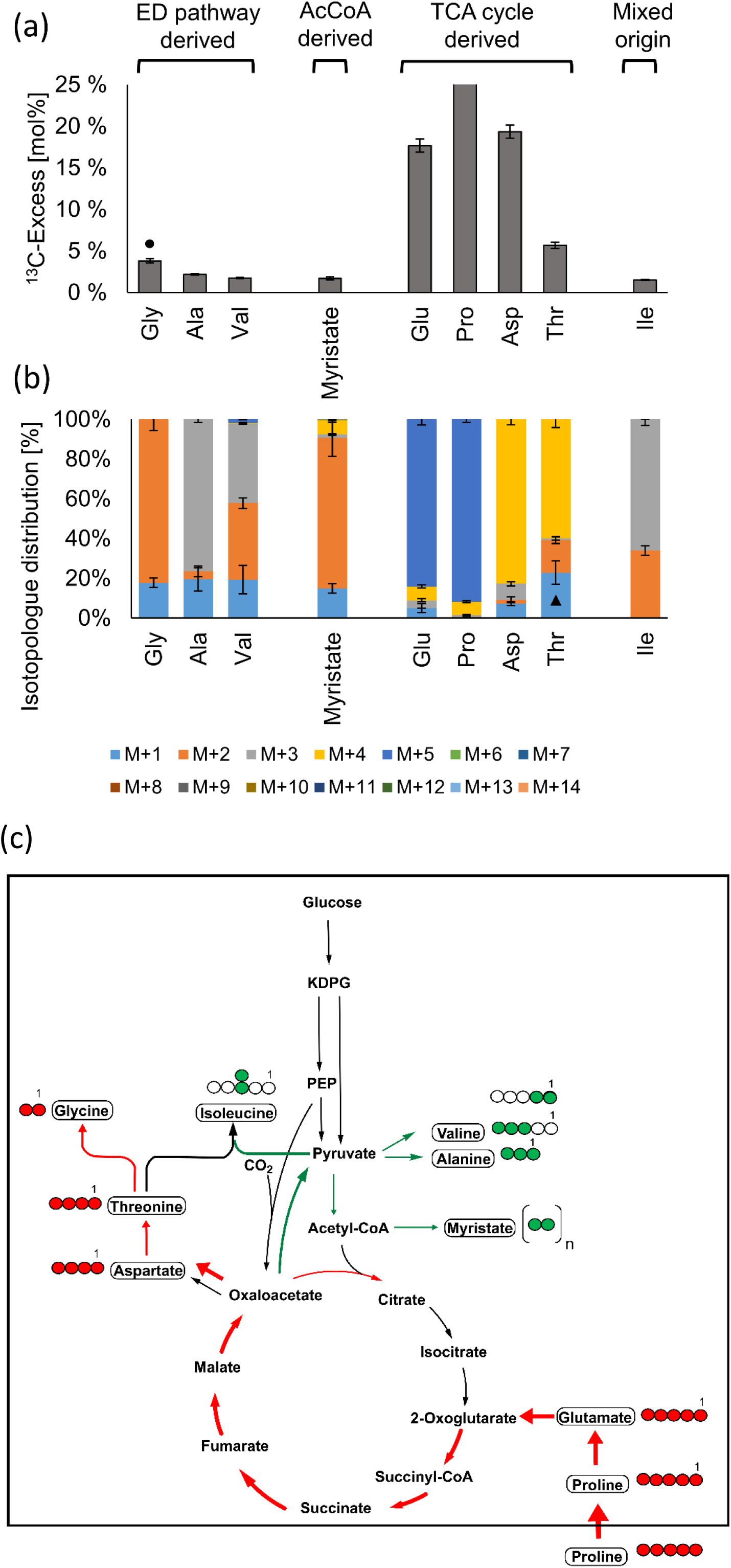
^13^C-Excess and isotopologue profiles of myristate and protein-derived amino acids from *N. gonorrhoeae* after growth in the presence of 0.4 mM [U-^13^C_5_]proline. (a) The ^13^C-excess (mol%) displays the overall ^13^C-incorporation in the respective metabolite beyond natural ^13^C-abundance. (b) Isotopologue compositions of myristate and amino acids with significant ^13^C-excess. M+1 from threonine might be overestimated due to the presence of another substance with the same retention time from the GC column and having the same mass as M+1 of threonine. c) Scheme of the carbon fluxes starting from the supplied [U-^13^C_5_]proline in the central carbon metabolism of *N. gonorrhoeae*. The red arrows indicate fluxes via conversion of proline into glutamate which feeds into 2-oxoglutarate and downstream intermediates of the TCA cycle. The green arrows indicate fluxes from oxaloacetate into pyruvate and amino acids thereof. Notably, no significant ^13^C-excess was detected for metabolites further upstream from pyruvate. For more details, see legend of Fig. 4.

**Fig. 6.**
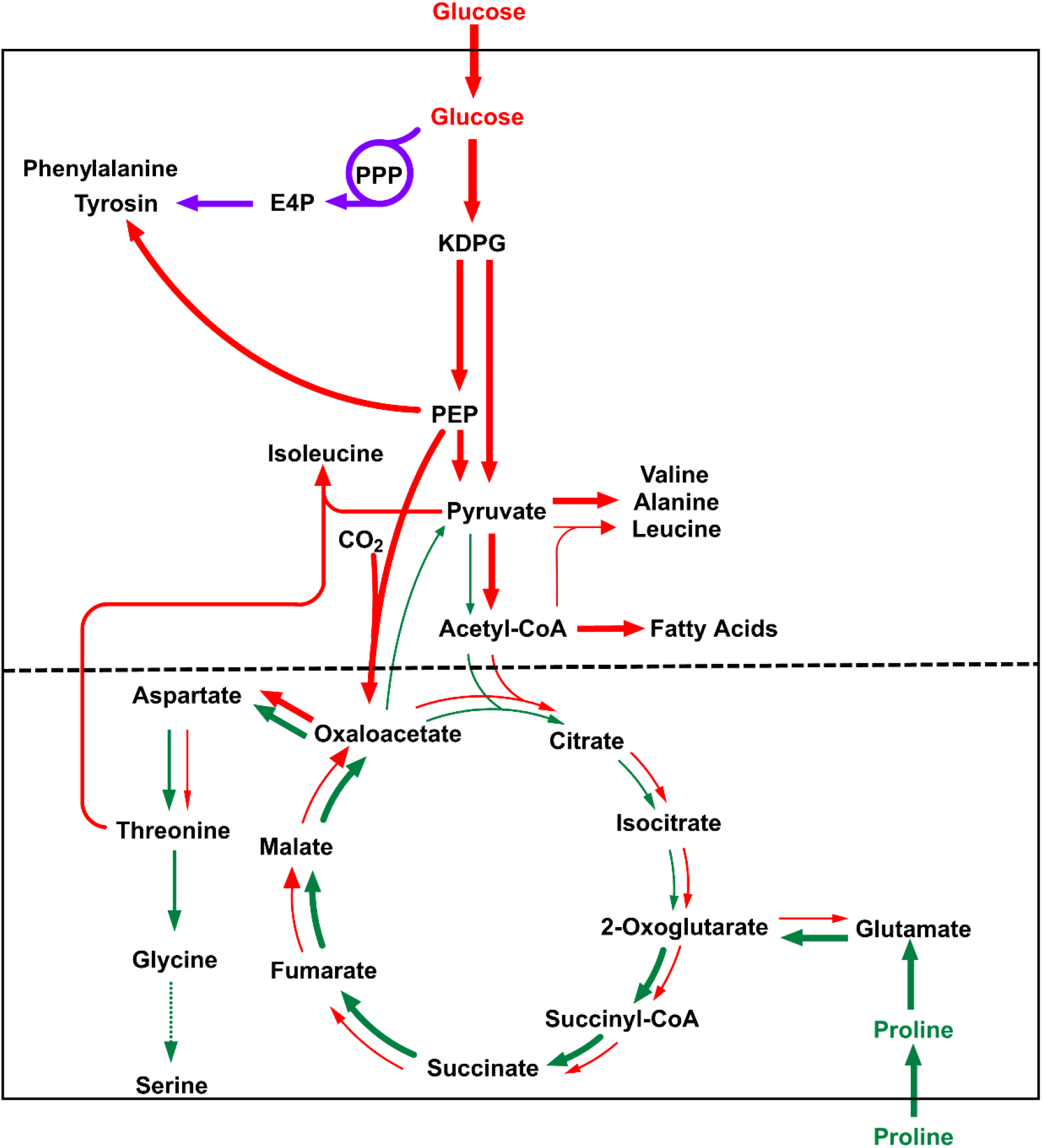
Model for the bipartite central carbon metabolism in *N. gonorrhoeae*. Red arrows indicate fluxes based on glucose utilization, purples arrows indicate fluxes via the PPP and the shikimate Pathway, and green arrows indicate fluxes determined by the usage of proline and related amino acids. The thickness of the arrows corresponds to the approximate relative extent of ^13^C-incorporation that was observed with the respective labelled tracers. The dashed black line indicates the borderline of the two metabolic modules in the bipartite metabolic network.

Based on this metabolic model, we now characterized the impact of the sibling sRNAs and targeted transport proteins on the core metabolism of *N. gonorrhoeae*. This was done by supplementing the respective mutant strains with [U-^13^C_6_]glucose or [U-^13^C_5_]proline during growth as explained above for the parent strain.

### NGFG_00045 and NGFG_01564 transport BCAAs and phenylalanine, respectively

The sibling sRNA targets NGFG_00045 and NGFG_01564 presumably encode amino acid transporters (Table 1, Fig. 3). Indeed, ^13^C-excess of alanine, isoleucine, leucine and valine differed significantly in the *N. gonorrhoeae* _Δ_45 mutant in comparison to the wild-type during growth with [U-^13^C_6_]glucose (Fig. 7a). While ^13^C-excess increased massively in the BCAAs isoleucine, leucine and valine (+16-35%), it decreased in alanine (−9%). The same effect, albeit to a much lower extent, was observed during growth of *N. gonorrhoeae* _Δ_45 with [U-^13^C_5_]proline (Fig. 7b). Here, also other amino acids showed minor differences, but the most significant effect was observed in glutamate and aspartate with increased ^13^C-excess values of 8-9% in the mutant. These complementary effects observed with different labelled substrates again corroborated the validity of the bipartite metabolic model introduced above (Fig. 6). Based on these observations, NGFG_00045 encodes a BCAA-transporter as *de novo* synthesis of this class of amino acids especially from [U-^13^C_6_]glucose was highly increased in the _Δ_45 mutant strain. All three of these amino acids use pyruvate as a building block (cf. Fig. 6); their increased biosynthesis depleted the pool of pyruvate available for alanine synthesis, and thereby reduced the ^13^C-excess in this amino acid. On the other hand, the synthesis of aspartate and glutamate was potentially upregulated to meet the increasing demand for aspartate as precursor in isoleucine biosynthesis (cf. Fig. 6) and for nitrogen donors in the transamination step of BCAA biosynthesis (Fig. 7c).

**Fig. 7.**
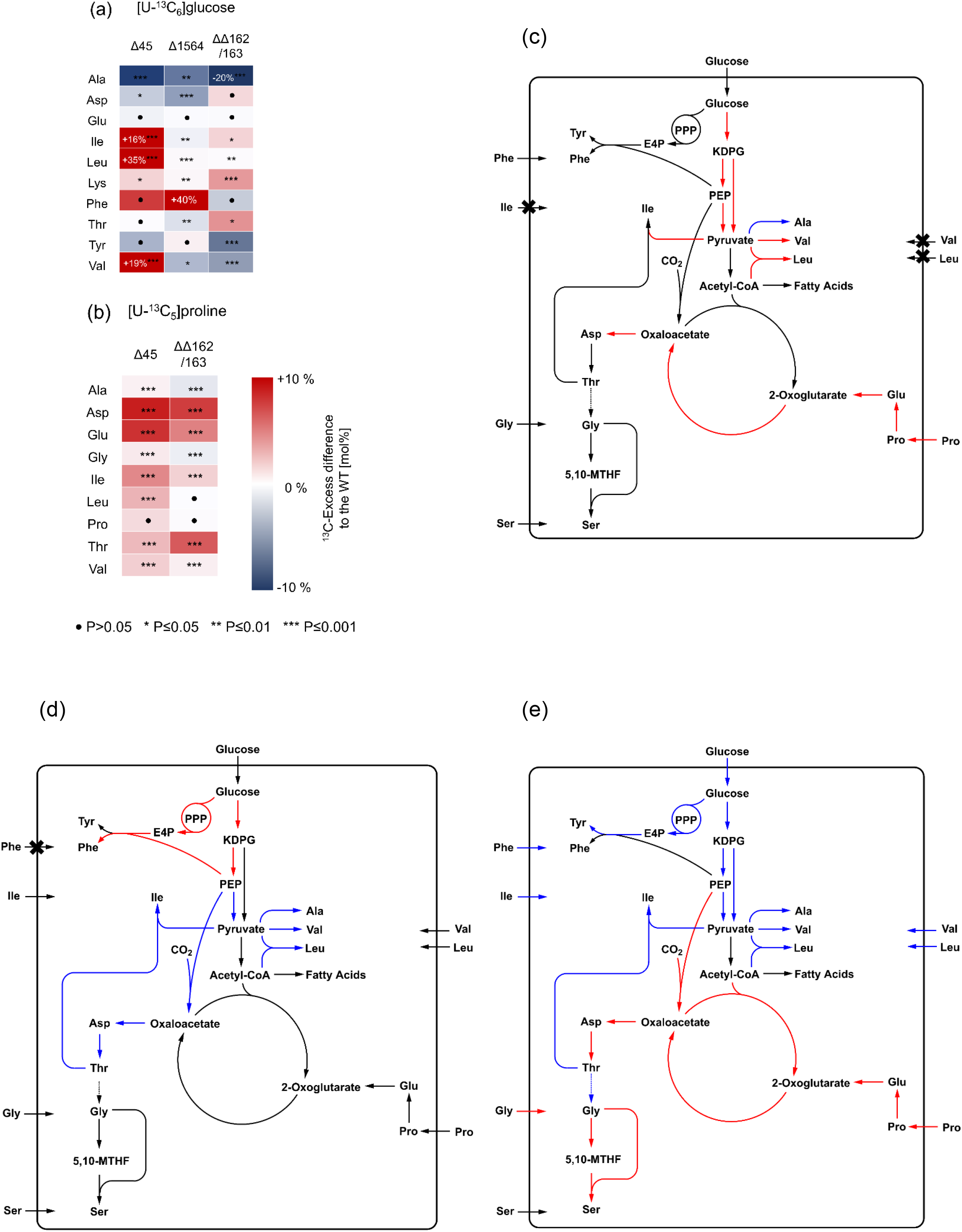
Differential labelling patterns and fluxes into protein-derived amino acids of *N. gonorrhoeae* MS11 (wild-type) and several mutant strains. (a) Heat map of ^13^C-excess from [U-^13^C_6_]glucose in amino acids from *N. gonorrhoeae* _ΔΔ_162/163, _Δ_45 and _Δ_1564 relative to the wild-type. (b) Heat map of ^13^C-excess from [U-^13^C_5_]proline in amino acids from *N. gonorrhoeae* _ΔΔ_162/163 and _Δ_45 relative to the wild-type. Numbers in the boxes indicate values outside of the color scale. Statistical significance was determined using Student’s t-test analysis (filled circles=not significant, *=p<0.05, **=p<0.01, ***=p<0.001). (c) Diffential fluxes in *N. gonorrhoeae* _Δ_45 relative to the wild-type strain. Red arrows indicate up-regulated fluxes, while blue arrows indicate down-regulated fluxes. The crosses indicate downregulated uptake of amino acids in the mutant. (d) Diffential fluxes in *N. gonorrhoeae* _Δ_1564 relative to the wild-type strain. Red arrows indicate up-regulated fluxes, while blue arrows indicate down-regulated fluxes. The crosses indicate downregulated uptake of phenylalanine in the mutant. (e) Diffential fluxes in the *N. gonorrhoeae* _ΔΔ_162/163 double mutant strain relative to the wild-type strain. Red arrows indicate up-regulated fluxes, while blue arrows indicate down-regulated fluxes and uptake of amino acids.

In case of the NGFG_01564 mutant (_Δ_1564), the labelling experiment with [U-^13^C_6_]glucose produced a huge increase in the ^13^C-excess of phenylalanine (+40%) when compared to the wild-type (Fig. 7a). Other significant differences (>1%) included a decrease in ^13^C-excess of alanine, valine, aspartate and threonine (−6.2 to −1.6%). These effects could again be explained by a decreased pool of labelled PEP and pyruvate as ^13^C-PEP was consumed for *de novo* phenylalanine biosynthesis in the _Δ_1564 mutant strain. As explained earlier, synthesis of aspartate from glucose heavily depends on the PEP carboxylase reaction in *N. gonorrhoeae* and aspartate is a precursor for threonine biosynthesis (cf. Fig. 6). Surprisingly, the ^13^C content of tyrosine was not significantly altered in the _Δ_1564 mutant strain (Fig. 7a), despite originating from the same metabolic precursors as phenylalanine. Based on these observations, NGFG_01564 encodes a phenylalanine transporter (Fig. 7d).

The identification of a previously non-characterized BCAA- and phenylalanine transporter as targets of the sibling sRNAs demonstrates the straightforward approach of our strategy to include ^13^C-based metabolomics as a tool to functionally analyse bacterial regulatory RNAs.

### Sibling sRNAs interfere with the TCA cycle, amino acid transport and metabolism

Looking at the sibling sRNA double-mutant strain *N. gonorrhoeae* _ΔΔ_162/163, a variety of effects could be observed in the labelling experiments. When supplementing [U-^13^C6]glucose (Fig. 7a), the ^13^C-excess in alanine, tyrosine and valine decreased (−5 to −20%), while the ^13^C-excess in isoleucine, lysine and threonine increased (+1.7 to +4%). In the experiment with [U-^13^C_5_]proline, the double-mutant strain showed an increased ^13^C-excess in aspartate, glutamate and again threonine and isoleucine (+5 to +7.6%). Alanine and glycine showed a decrease in ^13^C-excess of 1.1% and 0.7%, respectively (Fig. 7b).

As suggested earlier, the sibling sRNAs down-regulate the TCA cycle activity via negative control of various enzymes in the cycle (Bauer et al., 2017; Pannekoek et al., 2017). Consequently, in the double-mutant strain the TCA cycle was more active and utilized the supplied ^13^C-proline tracer at higher rates, thereby producing higher ^13^C-excess values in aspartate and glutamate, which are derived from oxaloacetate and proline, respectively (cf. Fig. 6). By closer examination of the isotopologue profiles, this effect can be distinguished from concomitant increased proline metabolization as a consequence of decreased BCAA uptake due to downregulation of NGFG_00045 (Fig. S4).

In the experiment with ^13^C-glucose, utilization of this substrate via the ED pathway as well as the PPP seems to be decreased leading to lower ^13^C-excess values in tyrosine and alanine. However, there was no significant effect observed in phenylalanine. As the sibling sRNAs upregulate the phenylalanine transporter NGFG_01564, there was less import of this amino acid in the double-mutant strain.

Therefore, the ^13^C-excess was much less diluted by import of exogenous phenylalanine compared to the wild-type. The same case can be made for the BCAAs valine, leucine and isoleucine, which were much less affected in the double-mutant strain compared to alanine, although they are also fully or partially derived from pyruvate. Herein, the sibling sRNAs upregulated the BCAA-transporter NGFG_00045, and consequently reduced import was observed in the double-mutant strain. However, genes in BCAA biosynthesis were also downregulated in mutant ΔΔ162/163 (Table 1), which could explain the lower ^13^C-excess in valine from ^13^C-glucose. Reduced glucose usage could be due to reduced glucose uptake, since mRNA levels of the putative glucose transporter NGFG_02263 were reduced in the sibling sRNA mutant (Table 1). Furthermore, increased uptake of unlabelled alanine by the sRNA double-mutant due to the upregulation of NGFG_01721 (Fig. 3) is likely to contribute to reduced ^13^C-accumulation in alanine.

Threonine presents an interesting case; when ^13^C-proline was supplemented, the ^13^C-excess of threonine increased but to smaller extent than its metabolic precursor aspartate. However, when ^13^C-glucose was used as a substrate, threonine showed a greater increase in ^13^C-excess than aspartate. Increased ^13^C-excess might be related to threonine acting as a precursor for glycine, which can be converted to serine by GlyA (reviewed in Stauffer, 2004). Increased uptake of unlabelled glycine in mutant _ΔΔ_162/163 (due to the upregulation of NGFG_01721) and elevated *glyA* expression resulted in an increased rate of serine production. These effects lead to a decreased ^13^C-excess in glycine and also mitigated the need for threonine as a precursor for glycine/serine thereby leading to the accumulation of labelled threonine in the double-mutant strain. This is in accordance with the labelling experiments, as in the double-mutant strain the ^13^C-excess was significantly increased in threonine, while ^13^C-excess in glycine was significantly decreased when supplementing [U-^13^C_5_]proline (Fig. 7b). The metabolic effects of sibling sRNA deletion are summerized in Fig. 7e.

## Discussion

The functional analysis of regulatory RNAs is hampered by the fact that often large targetomes are governed by them. Target identification by *in silico* predictions, quantification of mRNA or protein abundance in the presence or absence of the sRNA or RNA-seq based approaches exploiting direct sRNA-target mRNA interactions frequently yield numerous candidates which are then used to predict the function of the sRNA. Since sRNA-mediated modifications of mRNA or protein abundances are often moderate and metabolic pathways are frequently controlled by posttranslational modifications rather than protein abundance, very little is known about the true regulatory impact of the rapidly growing number of sRNAs with predicted metabolic targets (Sharma et al., 2007; Sheehan & Caswell, 2018; Gulliver et al., 2018; Potts et al. 2017; Beisel & Storz, 2011; Bækkedal & Haugen, 2015). We performed here a combined RNA-seq and metabolomics approach and thereby significantly extended the regulon controlled by the *N. gonorrhoeae* sibling sRNAs NgncR_162/163.

Our metabolomics analyses focussed on differentially transported metabolites (amino acids) leading to identification of function of thus far uncharacterized NgncR_162/163 targets like the potential BCAA-and phenylalanine transporters (NGFG_00045 and NGFG_01564, respectively). Metabolic flux analysis using isotopologue profiling confirmed the role of identified targets in metabolic pathways like the TCA and unveiled their role in orchestrating the substrate use in a newly identified bipartite metabolic network. The combination of transcriptome and carbon flux analysis followed here thus provides an excellent approach towards a real understanding of the outcome of riboregulation targeting metabolic processes in bacteria.

New members of the NgncR_162/163 targetome predominantly encode proteins involved in nutrient uptake and serine/glycine metabolism. This is reminiscent of *E. coli*/*Salmonella typhimurium* GcvB, which is conserved in members of the γ-proteobacteria (Sharma et al., 2007), and directly controls more than 50 mRNA targets comprising mostly periplasmic amino acid binding ABC transporter components, amino acid permeases (among others the glycine permease *cycA*) and enzymes involved in amino acid metabolism (Sharma et al., 2011; Lalaouna et al., 2019; Miyakoshi et al., 2021). GcvB is abundant during exponential growth in nutrient rich medium, but hardly detectable in bacteria from stationary phase or upon culture in minimal medium (Argaman et al., 2001; Sharma et al., 2007; Pulvermacher et al., 2009). The growth phase dependent expression pattern of GcvB results from the accumulation of the GcvB sponge SroC in stationary phase which triggers the RNase E-dependent degradation of GcvB via base-pairing interaction (Miyakoshi et al., 2015; Lalaouna et al., 2019). In addition, expression of GcvB is under control of GcvA, the transcriptional regulator of the glycine cleavage operon *gcvTHP* (Urbanowski et al., 2000) and is induced when glycine is available. Similar to GcvB, the *Neisseria* sibling sRNAs are most abundant during exponential growth under nutrient rich conditions, while sRNA levels decline in stationary phase (Fig. S5). Interestingly, in Rhizobiales, a plethora of ABC transport systems is regulated by sibling sRNAs (AbcR1 and AbcR2; reviewed in Sheehan & Caswell, 2018).

Newly identified targets are both under negative and positive control indicating that the sibling sRNAs not only act by obstructing the RBS as demonstrated previously (Bauer et al., 2017; Pannekoek et al., 2017; Heidrich et al., 2017), but also employ other means of target regulation triggered by binding of the mRNA within the CDS (Fig. 2; Fig. S1). *In silico* predicted regions of complementarity between the newly identified target mRNAs and the *N. gonorrhoeae* sibling sRNAs engage sequence motifs which are shared by both sRNA molecules (Fig. S1). However, in contrast to the well-established targets *prpB, prpC, ack* and NGFG_01721, full post-transcriptional regulation of which requires only one sibling (Bauer et al., 2017; Helmreich & Beier, unpublished), complete functional redundancy is not observed for the new members of the NgncR_162/163 regulon. Single complementation of MS11 ΔΔ162/163 under steady state conditions revealed a higher impact of NgncR_163 on target regulation (Fig. 1b), which might be explained by the higher abundance of NgncR_163 (Bauer et al., 2017). Such a diverse regulatory impact of siblings that differ in their abundance, but exhibit the same base pairing capability for the target has also been observed in case of post-transcriptional regulation of the salmochelin siderophor receptor IroN by *S. typhimurium* RhyB1 and RhyB2 (Balbontin et al., 2016). Our findings therefore suggest hierarchical target control based on the abundance of the *Neisseria* sibling sRNAs. However, besides growth phase-dependent differences in sRNA abundancy, which similarly apply to both siblings (Fig. S5), environmental conditions affecting expression of NgncR_162/163 could not be identified yet.

From isotopologue profiling experiments, we deduce a bipartite metabolism in *N. gonorrhoeae* where energy metabolism is mainly driven via amino acids like glutamate and proline which feed the TCA cycle, while glucose degradation via the ED and PPP pathway mostly provides intermediates for anabolic pathways (Fig. 6). This model is well in line with the observation that glutamate, glutamine and proline (and also asparagine), are consumed to the highest extent when gonococci are grown in chemically defined media (Fig. S2). Similar models for a bipartite metabolic network were already presented for other pathogenic bacteria such as *L. monocytogenes, Legionella pneumophila, Coxiella burnettii, Chlamydia trachomatis* and *Helicobacter pylori* (Grubmüller et al., 2014; Häuslein et al., 2016; Häuslein et al., 2017a; Mehlitz et al., 2017; Steiner et al., 2021). Downregulation of the citrate synthase GltA (and the citrate transporter NGFG_00249) by the sibling sRNAs (Bauer et al., 2017; Pannekoek et al., 2017) might promote the channeling of glutamate-derived 2-oxoglutarate into the TCA cycle. It is interesting to note that *gltA* is also a target of *Pasteurella multocida* GcvB (Gulliver et al., 2018). In the sRNA double-mutant, activity of the ED and PPP pathways was seemingly dampened, while the TCA cycle was more active, which is in accordance with the fact that several TCA cycle genes are under negative control of the sibling sRNAs (Table 1) (Bauer et al., 2017; Pannekoek et al., 2017). Interestingly, expression of the ED pathway enzymes glucose 6-phosphate 1-dehydrogenase (*zwf*) and 6-phosphogluconolactonase (*pgl*) in meningococci was reported to be negatively regulated by the transcription factor GdhR (Monaco et al., 2006), which itself is under negative control of the sibling sRNAs in *N. gonorrhoeae* (Bauer et al., 2017). Furthermore, in meningococci it was shown that ED pathway genes are upregulated in the presence of glucose, while TCA cycle genes and *gdhR* are downregulated (Antunes et al., 2016). However, *zwf* and *pgl* were not differentially expressed in a *gdhR* mutant of *N. gonorrhoeae* (Ayala & Shafer, 2019) and, consistently, we did not detect changes in *zwf* mRNA amounts in mutant ΔΔ162/163 by qRT-PCR (data not shown). Nevertheless, the labelling experiments clearly support the notion that the two putative modules of the bipartite metabolic network in *N. gonorrhoeae* are under opposite regulation by the sRNAs (Fig. 7e). This suggests a role for the sRNAs in optimizing growth of the pathogen while adapting to different environments during infection.

In this study, we demonstrate that the sibling sRNAs modulate BCAA (NGFG_00045)/phenylalanine (NGFG_01564) and glycine/alanine (NGFG_01721) import in a reciprocal manner (Fig. 3, Fig. 7). Derepression of glycine import in the absence of NgncR_162/163 is concomitant with upregulation of *gcvH* and *glyA* (Fig. 1a, Fig. S3, Fig. 7e), suggesting an important role of the sibling sRNAs in the regulation of serine-glycine metabolism, which in turn impacts on the biosynthesis of nucleotides, vitamins and other amino acids via the supply of C1-units. In fact, based on isotopologue profiling data, we propose that the threonine utilization (Tut) cycle (Ravnikar & Sommerville, 1986; Stauffer, 2004) is active in *Neisseria* to enable serine biosynthesis from threonine in the absence of 3-phosphoglycerate dehydrogenase which is not encoded in the genome of the pathogenic *Neisseria*. Both serine and threonine were efficiently taken up by wild-type gonococci during growth in chemically defined medium, while glycine concentration remained almost unchanged (Fig. S2). In contrast, in the sRNA double-mutant the need of serine uptake was compensated to a certain extent by an increased glycine uptake and cleavage via GcvTHP followed by a rise in serine biosynthesis from glycine and 5,10-MTHF due to upregulation of GlyA. This was accompanied by increased ^13^C-enrichment in threonine which was converted to glycine to a lesser extent (Fig. 7). Threonine being a precursor of serine might even give a hint to the biological relevance of positive regulation of the BCAA transporter NGFG_00045 by the sibling sRNAs: since threonine is also a precursor in the biosynthesis of isoleucine, increasing the uptake of isoleucine might spare threonine for use in serine synthesis. This hypothesis is further supported by the observation that compared to wild-type MS11 growth of mutant ΔΔ162/163 was less attenuated in chemically defined medium lacking serine (data not shown).

Expression of lactoylglutathione lyase, also named glyoxalase I (GloA) is positively regulated by the sibling sRNAs (Table 1; Fig. 1a). The glyoxalase system mediates the detoxification of the highly reactive compound methylglyoxal, which is a by-product of glycolysis and gluconeogenesis during the conversion of triose phosphate isomers. First, methylglyoxal and gluthathion are converted to S-lactoylglutathione by glyoxalase I, which is then cleaved by glyoxalase II yielding *D*-lactate and gluthathione (reviewed in Morgenstern et al., 2020). Interestingly, methylglyoxal is also formed during the catabolism of threonine via threonine dehydrogenase in the Tut-cycle, due to decarboxylation of α-amino-β-ketobutyrate and subsequent oxidation of the intermediate aminoacetone (Dutra et al., 2001). Upregulation of glyoxalase I is concomitant with downregulation of glycine uptake by the sibling sRNAs which we propose to result in increased Tut cycle activity. Other new members of the NgncR_162/163 regulon are related to the synthesis and maintenance of iron sulfur (Fe-S) centers (Table 1; Fig. 1a). DnrN is supposed to be involved in the repair of Fe-S centers damaged by oxidative or nitrosative stress (Overton et al., 2008). Interestingly, *dnrN* and two other positively regulated NgncR_162/163 targets, *norB* and *aniA*, were shown to be under control of the same transcription factor, NsrR (Overton et al., 2006). IscR is the transcriptional regulator of the *iscRSUA* operon encoding enzymes for Fe-S cluster biosynthesis. IscR itself contains an [Fe2-S2] cluster and holo-IscR was shown to directly repress expression of the *iscRSUA* operon in *E. coli* and pathogenic bacteria in order to maintain proper Fe-S cluster homeostasis (reviewed in Miller & Auerbuch, 2015). Both *dnrN* and *iscR* were deregulated in a Δ*hfq* mutant of *N. meningitidi*s (Fantappié et al., 2011) indicating post-transcriptional regulation. Since *dnrN* and *iscR* are inversely regulated by the sibling sRNAs with *dnrN* being activated and *iscR* being repressed, metabolic enzymes containing Fe-S clusters might be of particular importance under conditions when the sibling sRNAs are abundant. Besides glucose and pyruvate gonococci can use lactate as carbon and energy source, as electrons from the oxidation of both *L*- and *D*-lactate feed directly into the respiratory chain (Atack et al., 2014). Interestingly, one of the two gonococcal *L*-lactate dehydrogenases (LutACB) contains Fe-S clusters (Chen et al., 2020; Thomas et al., 2011) and lactate uptake is indirectly controlled by the sibling sRNAs via GdhR being a repressor of the *lctP* gene encoding lactate permease (Ayala & Shafer, 2019). Lactate permease is considered as a virulence factor of *N. gonorrhoeae*, since *lctP* deficient mutants are attenuated in a murine model of lower genital tract infection (Exley et al., 2011). It should be noted that RNA-seq also suggested positive regulation of NGFG_02263, encoding the ortholog of the menincococcal sole glucose transporter (Derkaoui et al., 2016), by the sibling sRNAs. In *N. meningitidis*, deletion of *hfq* resulted in downregulation of the glucose transporter transcript arguing in favour of direct or indirect sRNA-mediated expression control (Fantappié et al., 2011). However, the putative target NGFG_02263 could not be validated by qRT-PCR.

In conclusion, the work presented here expands our knowledge about the mechanisms of action of the sibling sRNAs NgncR_162/163 and their regulon. The data demonstrate that the siblings do not exhibit a complete functional redundancy and that they can act both as negative and positive regulators thus applying different mechanisms of action which, however, need to be characterized in more detail in the future. Moreover, the combined results of RNA-seq analysis and isotopologue profiling point to the operation of a bipartite central carbon metabolism in *N. gonorrhoeae* and a role of the sibling sRNAs in the regulatory networks which govern these central metabolic pathways and link them to the requirement of nutrient, and in particular, amino acid uptake. Thus, the sibling sRNAs appear to play a superior role in the regulatory hierarchy of central metabolic pathways of the gonococcus.

## Experimental Procedures

### Bacterial strains and growth conditions

The *N. gonorrhoeae* mutants used in this study were derived from wild-type strain MS11 (GenBank accession number NC_022240.1) and are listed in Table S1. *N. gonorrhoeae* was grown on GC agar (Oxoid) plates with 1% vitamin mix (Bauer et al., 2017) for 14-16 h at 37°C in a humidified 5% CO_2_ atmosphere. Liquid cultures were grown in PPM (proteose peptone #3 (15 g), soluble starch (1 g), KH_2_PO_4_ (4 g), K_2_HPO_4_ (1 g), NaCl (5 g/l dH_2_O) containing 1% vitamin mix and 0.04% (w/v) NaHCO_3_. Growth in chemically defined medium was conducted in CDM10 (Dyer et al., 1987) with slight modifications (*L*-glutamate: 0.0445 g/l; *L*-aspartate: 0.02 g/l). For metabolic labelling experiments *L*-proline and glucose were replaced by [U-^13^C_5_]L-proline (0.05 g/l; 0.4 mM) and [U- ^13^C_6_]D-glucose (2.5 g/l; 13,9 mM), respectively. Bacteria were grown to an OD_550_ = 0.6 for4 h in order to achieve steady state conditions in protein-derived amino acids and fatty acids which were subjected to isotopologue profiling. Media were supplemented with kanamycin, erythromycin or spectinomycin at final concentrations of 40 µg/ml, 7 µg/ml and 50 µg/ml, respectively, when required. *Escherichia coli* TOP10 (Thermo Fisher Scientific) and *E. coli* DH5α (Hanahan, 1983) were cultured in lysogeny broth (LB). When required, antibiotics were added to the following final concentrations: ampicillin, 100 µg/ml, kanamycin 30 µg/ml, chloramphenicol, 30 µg/ml.

### Construction of *N. gonorrhoeae* mutants

PCR primers for the amplification of DNA-fragments used for mutant construction are listed in Table S2. For the synthesis of *N. gonorrhoeae*-specific fragments chromosomal DNA of strain MS11 was used as template. Clonings were performed in *E. coli* DH5α.

MS11 P_*opa*_45, MS11 ΔΔP_*opa*_45 and MS11 ΔΔP_*opa*_45c: A 648 bp DNA-fragment from the upstream region of NGFG_00045 was amplified using primer pair 45-5UTR-1/45-5UTR-23 and was cloned into vector plasmid pSL1180 (Brosius, 1989) together with an erythromycin resistance cassette amplified with primers 45-5ermC-13/45-5ermC-23 from plasmid pMR68 (Ramsey et al., 2012). The combined DNA-segments were then amplified with the outer primers 45-5UTR-1/ermPopa1. The *opa* promoter was PCR-amplified with primer pair ermPopa2/45Popa-4 and was combined with a DNA-fragment comprising the 5’-UTR (Remmele et al., 2014) and 546 bp of the coding region of NGFG_00045 (amplified with primer pair 45Popa-5/45Flag-6) by overlap extension PCR. Finally, the DNA-segments consisting of the upstream region of NGFG_00045 and *ermC* and the P_*opa*_-NGFG_00045 fusion were combined by overlap extension PCR (using primers 45-5UTR-1/45Flag-6) and the resulting DNA-fragment was transformed into *N. gonorrhoea* MS11 and ΔΔ162/163 (Bauer et al., 2017) to yield strains P_*opa*_45 and ΔΔP_*opa*_45. For complementation ΔΔP_*opa*_45 was transformed with plasmid pMR-162/163 (Bauer et al., 2017).

MS11 P_45_gfp and MS11 ΔΔP_45_gfp: The fusion of the upstream region of NGFG_00045, including the 5’-UTR, to the *gfp*-mut2 gene was constructed by combination of DNA-fragments amplified with primer pairs 45gfp-1/45gfp-3 and 45gfp-2/45gfp-7 via overlap extension PCR. Plasmid pKEN (Cormack et al., 1996) was used as template for the amplification of *gfp*-mut2. The resulting DNA-fragment was combined with a DNA-fragment comprising *ermC* and 500 bp from the downstream region of NGFG_00045, which was obtained by overlap extension PCR using DNA-fragments amplified with primer pairs 45-5ermC-13/45-3ermC-23 and 45gfp-6/45mut-5, respectively. Transformation of the resulting DNA-fragment into *N. gonorrhoeae* MS11 and ΔΔ162/163 yielded strains P_45_gfp and ΔΔP_45_gfp in which NGFG_00045 is replaced by *gfp*. A consensus Shine-Dalgarno sequence was introduced in the NGFG_00045 5’UTR by performing overlap extension PCR using DNA-fragments amplified with primer pairs 45gfp-1/45gfp-8 and 45gfp-9/45mut-5 from chromosomal DNA of strain P_45_gfp. The combined DNA-fragment was transformed into *N. gonorrhoeae* MS11 and ΔΔ162/163 to yield strains P_45_gfpSD and ΔΔP_45_gfpSD.

MS11 Δ45 and MS11 Δc45: To create mutant Δ45 a DNA-segment covering 289 bp from the upstream region and the sequence encoding the first 80 amino acids of NGFG_00045 was replaced by an erythromycin resistance cassette via allelic exchange mutagenesis. In the DNA-fragment used for transformation of MS11 the *ermC* gene (amplified with primer pair 45-5ermC-13/45-3ermC-3) is flanked by sequences comprising the 3’-end of NGFG_00044 and part of the intergenic region between NGFG_00044 and NGFG_00045 (amplified with primer pair 45-5UTR-1/45-5UTR-23) and encoding amino acids 81 to 239 of NGFG_00045 (amplified with primer pair D451/D45-2). For complementation overlap extension PCR was applied to insert a kanamycin resistance cassette (amplified with primer pair D45-4/D45-5) between the upstream fragment used for construction of Δ45 (amplified with primer pair 45-5UTR-1/D45-3) and a DNA-segment covering promoter region, 5’-UTR and 785 bp from the 5’-end of NGFG_00045 (amplified with primer pair D45-5/D45-6). The resulting DNA-fragment was transformed into Δ45 to yield Δc45.

MS11 Δ1564: In mutant Δ1564 the region encoding amino acids 1 - 243 of NGFG_01564 is replaced by an erythromycin resistance cassette. A DNA-fragment comprising segments derived from the upstream (PCR amplified with primer pair D1564-1/D1564-2) and coding region of NGFG_01564 (covering amino acid 245 to 412, amplified with primer pair D1564-5/D1564-6) which flank the *ermC* gene (amplified with primer pair D1564-5/D1564-6) was assembled by overlap extension PCR and transformed into MS11. Homologous recombination yielded strain Δ1564.

MS11 Δ1721: In mutant Δ1721 ORF NGFG_01721 as well as 248 bp from its upstream region are substituted by a spectinomycin resistance cassette. To construct this mutant a SacI/PstI fragment comprising part of the upstream ORF NGFG_01720 and intergenic region was amplified with primer pair 1721up1/1721up2 and was cloned together with a spectinomycin resistance cassette into pSL1180. The spectinomycin cassette expressing *aadA1* under control of the *Neisseria* P_*opa*_ promoter was amplified from MS11 ΔtfpR2 (Zachary et al., 2021) using primer pair spec2(PstI)/Popa5(KpnI). The assembled fragments were subsequently amplified with primer pair 1721up1/D1721-4 and were combined via overlap extension PCR with a DNA-fragment derived from the downstream region of NGFG_01721 (amplified with primer pair D1721-3/D1721-2). Transformation of MS11 with the full-length DNA-fragment yielded strain Δ1721.

### RNA preparation, RNA-Seq and analysis of RNA-Seq data

*N. gonorrhoeae* MS11 and ΔΔ162/163 were grown to an OD_550_ = 0.5 in PPM. RNA was prepared using the miRNeasy Micro Kit (Qiagen) according to the manufacturer’s instructions followed by DNaseI treatment. RNA integrity was checked using a Bioanalyser. After enrichment of mRNA using the Universal Ribodepletion Kit cDNA preparation was performed with the Next Ultra Directional Library Preparation Kit for Illumina (NEB). The cDNA was sequenced on HiSeq 3000 (Illumina) yielding 100 bp paired end reads. Reads with a minimum length of 15 bp after removal of low quality ends and adapters using cutadapt (Martin, 2011), were mapped to the *N. gonorrhoeae* MS11 genome (Genome assembly: ASM15685v2; (Ribeiro et al., 2012). Read mapping was conducted using Bowtie2 (Langmead & Salzberg, 2012). Genes were quantified using featureCounts (Liao et al., 2014). DESeq2 (Love et al., 2014) was used to identify differentially regulated transcripts.

### Northern Blot analysis and real-time quantitative PCR

Northern Blot analysis was performed as described previously (Bauer et al., 2017). For qRT-PCR experiments 1 µg of RNase-free DNase-treated RNA was reverse transcribed with random hexamer primers using RevertAid first strand cDNA Synthesis Kit (Thermo Scientific). All qRT-PCR reactions were performed in triplicate in a 20 µl mixture containing cDNA (5 µl of 1:20 dilution), PerfeCTa SYBR Green FastMix containing ROX (Quanta Biosciences) and 18 pmol of primer (Table S2). Amplification and detection of PCR products was performed with a StepOne Plus qRT-PCR system (Applied Biosystems) using the following procedure: 95°C for 10 min, then 40 cycles of 95°C for 15 sec and 60°C for 60 sec followed by dissociation curve analysis. The relative expression levels of the genes studied were normalized to the 5S rRNA gene. Data were analysed using the ΔΔC_T_ method (Livak & Schmittgen, 2001). If not stated otherwise, at least three qRT-PCR experiments were performed in triplicate with cDNA which was reverse transcribed from independent RNA preparations.

### Immunoblot analysis

*N. gonorrhoeae* were grown to an OD_550_ = 0.6 in PPM. Cells from 1 ml of culture were harvested by centrifugation and resuspended in 50 µl of Laemmli buffer and incubated for 7 min at 95°C. Western blot analysis of the samples was performed as described previously (Bauer et al., 2017).

### Analysis of amino acid composition of (spent) culture media

Amino acids in bacterial culture medium were measured by UPLC-ESI–MS/MS using an Acquity Ultra Performance Chromatography system UPLC combined with a Quattro Premier triple quadrupole mass spectrometer (Waters, Milford, USA). Derivatization, chromatographic separation and successive detection of amino acids was carried out as described by Salazar et al. (Alterman & Hunziker, 2012) with modifications.

After sterile filtration, 100 µl of medium were diluted with 100 µl of methanol containing norvaline as internal standard, with a final concentration of 1mM. 20 µl of this mixture were used for subsequent amino acid derivatization using the AccQ-Tag Ultra Derivatization Kit (Waters, Milford, USA) following the manufacturer’s instructions.

Chromatographic separation was carried out using a BEH C18 column (2.1 × 100 mm, 1.7 µm particle size; Waters) equipped with a VanGuard pre-column and an in-line particle filter. Elution was performed with 100% eluent A for 1 min, followed by a binary solvent gradient to 30% of eluent B within 12 min at a flow rate of 0.4 ml/min. Eluent A consisted of 0.1% formic acid in water and eluent B of 0.1% formic acid in acetonitrile.

The ESI source was operated in positive mode at a source temperature 120°C with a capillary voltage set to 3 kV, the cone voltage at 30 V and the desolvation gas at 850 l/h at 400°C. Compounds were detected by multiple reaction monitoring (MRM, [M+H]^+^ _→_ m/z 171) with a dwell time of 25 ms and a collision energy of 18 V using argon as collision gas at a flow rate of 0.3 ml/min. Data acquisition and processing was carried out using MassLynx and QuanLynx (Waters, Milford, USA; version 4.1).

### Sample preparation for ^13^C-analysis of protein bound amino acids

The analysis of protein bound amino acids was done as previously described (Eylert et al., 2008). About 1 mg of lyophilized bacterial cell pellet was hydrolysed over night at 105°C after having added 500 µl HCl (6 M). The hydrolysate was dried under a gentle stream of nitrogen at 70°C and dissolved in 200 µl of acetic acid (50%). For the isolation of protein bound amino acids, a cation exchange column of Dowex 50WX8 (H^+^-form; 7×10 mm; 200-400 mesh, 34-74 µm) was washed with 1000 µl of methanol (70%) and 1000 µl of H_2_O (bidest.). After applying the sample, which was dissolved in acetic acid, to the column, the column was first evolved with 1600 µl of H_2_O (bidest.). Subsequently, the amino acids were eluted with 1000 µl of aqueous ammonia solution (4 M). After drying the ammonia eluate under a gentle stream of nitrogen at 70°C, the isolated amino acids were incubated with 50 µl *N*-methyl-*N*-*tert*-butyldimethylsilyltrifluoroacetamide (MTBSTFA) containing 1% *tert*-butyldimethylchlorsilane and 50 µl acetonitrile (anhydrous) for 30 min at 70°C. The *N*-*tert*-butyldimethylsilyl-derivates (TBDMS) of the amino acids were analysed by GC-MS.

### Sample preparation for ^13^C-analysis of fatty acids

The analysis of fatty acids was done as previously described (Häuslein et al., 2017a). In brief, about 5 mg of the lyophilized bacterial cell pellet was dissolved in 1 ml cold methanol and 800 mg of glass beads were added. Cells were mechanically disrupted and lysed using a ribolyser system (Hybaid) with three cycles of 20 s at 6.5 m s^-1^. Then, the samples were centrifuged for 10 min at 7000 rpm and the supernatant was subsequently dried under a gentle stream of nitrogen at room temperature. For derivatization, the dry residue was incubated with 50 µl *N*-methyl-*N*-*tert*-butyldimethylsilyltrifluoroacetamide (MTBSTFA) containing 1% *tert*-butyldimethylchlorsilan and 50 µl acetonitrile (anhydrous) for 1 h at 70°C. The *N*-*tert*-butyldimethylsilyl-metabolites (TBDMS) were analysed by GC-MS.

### GC-MS measurement parameters

For the analysis of TBDMS-amino acids, a QP2010 Plus gas chromatograph-mass spectrometer was used as previously described (Eylert et al., 2008). The column was heated to 150°C, kept at 150°C for 3 min, heated to 280°C with a temperature gradient of 7°C min^-1^ and kept at 280° for 3 min. For analysis of TBDMS-fatty acids (Häuslein et al., 2017a), the colum was kept at 100°C for 2 min and subsequently heated to 234°C (3°C min^-1^). Then the column was heated with 1°C min^-1^ to 237°C. Finally, the column was heated to 260°C (3°C min^-1^).

Each sample was measured in triplicate in order to account for technical errors. GC-MS data were processed with the Shimadzu LabSolution software V4.20. For the calculation of ^13^C-excess values and isotopologue profiles, Isotopo software was used (Ahmed et al., 2014).

## Acknowledgements

Roy Gross is acknowledged for critical reading of the manuscript. This work was funded by the Deutsche Forschungsgemeinschaft (DFG) grant RU 631/12-1 to TR and EI 384/16-1 to WE. The Core Unit was supported by the DFG – project numbers 179877739 and 316629583.

## Author Contributions

Design of the study: DB, WE, TR

Data aquisition: TS, SR, MZ, SB, BH, MKr, MJM

Data analysis and interpretation: TS, SR, WE, MZ, MKl, MKr, DB

Writing of the manuskript: DB, TS, WE, TR

## Data Availability Statement

RNA-seq data obtained in this study have been deposited with GEO under accession number GSE177032.

## Legends for Supplementary Figures

Fig. S1.

In silico prediction of sRNA-mRNA interactions. (a) Prediction of the secondary structure of NgncR_162 using the RNAfold web server (http://rna.tbi.univie.ac.at/cgi-bin/RNAWebSuite/RNAfold.cgi). Stem loops (SL) 1-3 and single stranded regions (SSR) 1 and 2 are indicated. Regions of complementarity between the sibling sRNAs and their target genes were analysed with IntaRNA (Mann et al., 2017) and were predicted within the 5’-UTR (b) or coding region (c). With the exception of NGFG_01553 and *gloA* the target-sRNA interactions shown apply to both NgncR_162 and NgncR_163 with only minor variations. Nucleotides which differ in the other sibling resulting in mismatches are underlined in the sequence of the depicted sRNA. In case of *gloA* complementarity to NgncR_163 is less pronounced and covers only nucleotides 204 - 226. Numbers refer to the nucleotide positions with respect to the translational start site (+1) in the case of mRNAs and the transcription initiation site in the case of NgncR_162 or NgncR_163. The start codon is marked in bold.

Fig. S2

Amino acid consumption by *N. gonorrhoeae* MS11 grown in a chemically defined medium. *N. gonorrhoeae* MS11 was grown to OD_550_ = 0.6 in modified CDM10 medium. Bars depict the amino acid concentration (mM) determined for CDM10 medium and spent medium from MS11 culture by mass spectrometry and represent the mean of four biological replicates (five technical replicates each). Error bars indicate the standard deviation.

Fig. S3

Post-transcriptional regulation of *glyA* by the sibling sRNAs NgncR_162/163. (a) GlyA transcript amounts were quantified by qRT-PCR in wild-type MS11, ΔΔ162/163 and the complemented strain ΔΔc162/163. The ratios of the transcript amount relative to the wild-type MS11 (normalized to 1) are depicted. Error bars indicate the standard deviation (n = 3). Statistical significance was determined using Student’s t-test analysis (**=p<0.01). (b) Expression of a translational *glyA*-*gfp* fusion was analysed in *E. coli* in the absence and presence of sRNA NgncR_162. *E. coli* Top10 were co-transformed with plasmid pXG-glyA (expressing a translational *glyA*-*gfp* fusion) and either plasmid pJV300 expressing a nonsense RNA (lane 1) or plasmids expressing NgncR_162 (pJV-162; lane 2) or a mutated derivative of NgncR_162 (pJV-162m1; lane 3). In pJV-162m1 the loop sequence of the NgncR_162 SL2 was changed from TTCTCCTT to TTCCAAGCT (Bauer et al., 2017). Equal amounts of protein were separated on a 12% polyacrylamide gel and Western blot analysis was performed with monoclonal antibodies directed against GFP and HSP60 used as loading control. The figure shows the results from a representative experiment (n = 2).

Fig. S4

Isotopologue profiles of glutamate (a) and aspartate (b) from wild-type MS11, _ΔΔ_162/163 and _Δ_45 after growth in CDM10 medium supplemented with 13.9 mM [U-^13^C_6_]glucose. Analysis of the isotopologue profiles of aspartate and glutamate allows for the differentiation of metabolic effects in ΔΔ162/163 and Δ45: In the _Δ_45 mutant, proline uptake is increased but expression of several TCA cycle enzymes is still repressed by the sibling sRNAs. When supplementing [U-^13^C_6_]glucose as a tracer, the abundance of heavier isotopologues (>M+2) in aspartate and glutamate decreased as the labelling in the TCA cycle was diluted due to the increased uptake of unlabelled proline in comparison to the wild-type. This lowered the probability for combination of two labelled precursors that would potentially produce heavier isotopologues in aspartate and glutamate. In the _ΔΔ_162/163 mutant strain, a similar effect albeit to a lower extent should be expected as the expression of NGFG_00045 is no longer upregulated by the sibling sRNAs. However, as the TCA cycle is also no longer repressed more labelled precursors entered the cycle and subsequently produced isotopologue profiles in aspartate and glutamate that were similar to the wild-type.

Fig. S5

The sibling sRNAs NgncR_162/163 are less abundant in stationary growth phase. (a) Northern Blot analysis was performed on RNA extracted from bacteria from logarithmic phase, the transition phase between logarithmic growth and stationary phase and stationary phase using radioactively labelled sRNA-specific oligonucleotides. Probing for 5S rRNA was used as loading control. The position of a radiolabelled marker RNA of 100 nucleotides is indicated on the left side of the panel. Images from four independent experiments were used for quantification of sRNA amounts. In (b) the growth curve of *N. gonorrhoeae* MS11 is shown and time points at which samples for RNA preparation were taken are indicated. (c) The mRNA abundance of target NGFG_01721, which is negatively regulated by the sibling sRNAs, was quantified by qRT-PCR in bacteria grown to logarithmic and stationary phase, respectively. qRT-PCR analysis was performed on cDNA from two independent RNA preparations.

## References

Ahmed Z, Zeeshan S, Huber C, Hensel M, Schomburg D, Münch R, Eylert E, Eisenreich W, Dandekar T. 2014. ‘Isotopo’ a database application for facile analysis and management of mass isotopomer data. Database (Oxford) 2014: bau077. https://doi.org/10.1093/database/bau077.

Alterman MA, Hunziker P (Eds). 2012. Amino Acid Analysis: Methods and Protocols. Humana Press. p 13–28. https://doi.org/10.1007/978-1-61779-445-2.

Antunes A, Golfieri, Ferlicca F, Giuliani MM, Scarlato V, Delany I. 2015. HexR controls glucose-responsive genes and central carbon metabolism in Neisseria meningitidis. J Bacteriol 198:644–654. https://doi.org/10.1128/JB.00659-15.

Argaman L, Hershberg R, Vogel J, Bejerano G, Wagner EG, Margalit H, Altuvia S. 2001. Novel small RNA-encoding genes in the intergenic regions of Escherichia coli. Curr Biol 11:941–950. https://doi.org/10.1016/S0960-9822(01)00270-6.

Atack JM, Ibranovic I, Ong CL, Djoko KY, Chen NH, Vanden Hoven R, Jennings MP, Edwards JL, McEwan AG. 2014. A role for lactate dehydrogenases in the survival of Neisseria gonorrhoeae in human polymorphonuclear leukocytes and cervical epithelial cells. J Infect Dis 210:1311–1318. https://doi.org/10.1093/infdis/jiu230.

Ayala JC, Shafer WM. 2019. Transcriptional regulation of a gonococcal gene encoding a virulence factor (L-lactate permease). PLoS Pathog 15:e1008233. https://doi.org/10.1371/journal.ppat.1008233.

Azam MS, Vanderpool CK. 2020. Translation inhibition from a distance: The small RNA SgrS silences a ribosomal protein S1-dependent enhancer. Mol Microbiol 114:391–408. https://doi.org/10.1111/mmi.14514.

Bækkedal C, Haugen P. 2015. The Spot 42 RNA: A regulatory small RNA wth roles in the central metabolism. RNA Biol 12:1071–1077. https://doi.org/10.1080/15476286.2015.1086867.

Balbontín R, Villagra N, Pardos de la Gándara M, Mora G, Figueroa-Bossi N, Bossi L. 2016. Expression of IroN, the salmochelin siderophore receptor, requires mRNA activation by RyhB small RNA homologues. Mol Microbiol 100:139–155. https://doi.org/10.1111/mmi.13307.

Bauer S, Helmreich J, Zachary M, Kaethner M, Heinrichs E, Rudel T, Beier D. 2017. The sibling sRNAs NgncR_162 and NgncR_163 of Neisseria gonorrhoeae participate in the expression control of metabolic, transport and regulatory proteins. Microbiology (Reading) 163:1720–1734. https://doi.org/10.1099/mic.0.000548.

Beisel CL, Storz G. 2011. The base-pairing RNA Spot 42 participates in a multioutput feedforward loop to help enact catabolite repression in Escherichia coli. Mol Cell 41:286–297. https://doi.org/10.1016/j.molcel.2010.12.027.

Bobrovskyy M, Vanderpool CK, Richards GR. 2015. Small RNAs regulate primary and secondary metabolism in Gram-negative bacteria. Microbiol Spectr 3. https://doi.org/10.1128/microbiolspec.MBP-0009-2014.

Brosius J. 1989. Superpolylinkers in cloning and expression vectors. DNA 8:759–777. https://doi.org/10.1089/dna.1989.8.759.

Caswell CC, Oglesby-Sherrouse AG, Murphy ER. 2014. Sibling rivalry: related bacterial small RNAs and their redundant and non-redundant roles. Front Cell Infect Microbiol 4:151. https://doi.org/10.3389/fcimb.2014.00151.

Chen NH, Ong CY, O’sullivan J, Ibranovic I, Davey K, Edwards JL, McEwan AG. 2020. Two distinct L-lactate dehydrogenases play a role in the survival of Neisseria gonorrhoeae in cervical epithelial cells. J Infect Dis 221:449–453. https://doi.org/10.1093/infdis/jiz468.

Chung GT, Yoo JS, Oh HB, Lee YS, Cha SH, Kim SJ, Yoo CK. 2008. Complete genome sequence of Neisseria gonorrhoeae NCCP11945. J Bacteriol 190:6035–6036. https://doi.org/10.1128/JB.00566-08.

Corcoran CP, Podkaminski D, Papenfort K, Urban JH, Hinton JC, Vogel J. 2012. Superfolder GFP reporters validate diverse new mRNA targets of the classic porin regulator, MicF RNA. Mol Microbiol 84:428–445. https://doi.org/10.1111/j.1365-2958.2012.08031.x.

Cormack BP, Valdivia RH, Falkow S. 1996. FACS-optimized mutants of the green fluorescent protein (GFP). Gene 173:33–38. https://doi.org/10.1016/0378-1119(95)00685-0.

Cox DL, Baugh CL. 1977. Carboxylation of phosphoenolpyruvate by extracts of Neisseria gonorrhoeae. J Bacteriol 129:202–206. https://doi.org/10.1128/jb.129.1.202-206.1977.

Darfeuille F, Unoson C, Vogel J, Wagner EG. 2007. An antisense RNA inhibits translation by competing with standby ribosomes. Mol Cell 26:381–392. https://doi.org/10.1016/j.molcel.2007.04.003.

Derkaoui M, Antunes A, Nait Abdallah J, Poncet S, Mazé A, Ma Pham QM, Mokhtari A, Deghmane AE, Joyet P, Taha MK, Deutscher J. 2016. Transport and catabolism of carbohydrates by Neisseria meningitidis. J Mol Microbiol Biotechnol 26:320–332. https://doi.org/10.1159/000447093.

Dutra F, Knudsen FS, Curi D, Bechara EJ. 2001. Aerobic oxidation of aminoacetone, a threonine catabolite: iron catalysis and coupled iron release from ferritin. Chem Res Toxicol 14:1323–1329. https://doi.org/10.1021/tx015526r.

Dyer DW, West EP, Sparling PF. 1987. Effects of serum carrier proteins on the growth of pathogenic neisseriae with heme-bound iron. Infect Immun 55:2171–2175. https://doi.org/10.1128/IAI.55.9.2171-2175.1987.

Exley RM, Wu H, Shaw J, Schneider MC, Smith H, Jerse AE, Tang CM. 2007. Lactate acquisition promotes successful colonization of the murine genital tract by Neisseria gonorrhoeae. Infect Immun 75:1318–24. https://doi.org/10.1128/IAI.01530-06.

Eylert E, Schär J, Mertins S, Stoll R, Bacher A, Goebel W, Eisenreich W. 2008. Carbon metabolism of Listeria monocytogenes growing inside macrophages. Mol Microbiol 69:1008–1017. https://doi.org/10.1111/j.1365-2958.2008.06337.x.

Fantappiè L, Oriente F, Muzzi A, Serruto D, Scarlato V, Delany I. 2011. A novel Hfq-dependent sRNA that is under FNR control and is synthesized in oxygen limitation in Neisseria meningitidis. Mol Microbiol 80:507–23. https://doi.org/10.1111/j.1365-2958.2011.07592.x.

Gripenland J, Netterling S, Loh E, Tiensuu T, Toledo-Arana A, Johansson J. 2010. RNAs: regulators of bacterial virulence. Nat Rev Microbiol 8:857–866. https://doi.org/10.1038/nrmicro2457.

Grubmüller S, Schauer K, Goebel W, Fuchs TM, Eisenreich W. 2014. Analysis of carbon substrates used by Listeria monocytogenes during growth in J774A.1 macrophages suggests a bipartite intracellular metabolism. Front Cell Infect Microbiol 3:156. https://doi.org/10.3389/fcimb.2014.00156.

Gulliver EL, Wright A, Lucas DD, Mégroz M, Kleifeld O, Schittenhelm RB, Powell DR, Seemann T, Bulitta JB, Harper M, Boyce JD. 2018. Determination of the small RNA GcvB regulon in the Gram-negative bacterial pathogen Pasteurella multocida and identification of the GcvB seed binding region. RNA 24:704–720. https://doi.org/10.1261/rna.063248.117.

Hanahan D. 1983. Studies on transformation of Escherichia coli with plasmids. J Mol Biol 166:557–580. https://doi.org/10.1016/s0022-2836(83)80284-8.

Häuslein I, Manske C. Goebel W, Eisenreich W, Hilbi H. 2016. Pathway analysis using ^13^C-glycerol and other carbon tracers reveals a bipartite metabolism of Legionella pneumophila. Mol Microbiol 100:229–246. https://doi.org/10.1111/mmi.13313.

Häuslein I, Cantet F, Reschke S., Chen F, Bonazzi M, Eisenreich W. 2017a. Multiple substrate usage of Coxiella burnetii to feed a bipartite metabolic network. Front Cell Infect Microbiol 29:285. https://doi.org/10.3389/fcimb.2017.00285.

Häuslein I, Sahr T, Escoll P, Klausner N, Eisenreich W, Buchrieser C. 2017b. Legionella pneumophila CsrA regulates a metabolic switch from amino acid to glycerolipid metabolism. Open Biol 7:170149. https://doi.org/10.1098/rsob.170149.

Hebeler BH, Morse SA. 1976. Physiology and metabolism of pathogenic neisseria: tricarboxylic acid cycle activity in Neisseria gonorrhoeae. J Bacteriol 128:192–201. https://doi.org/10.1128/jb.128.1.192-201.1976.

Heidrich N, Bauriedl S, Barquist L, Li L, Schoen C, Vogel J. 2017. The primary transcriptome of Neisseria meningitidis and its interaction with the RNA chaperone Hfq. Nucleic Acids Res 45:6147–6167. https://doi.org/10.1093/nar/gkx168.

Holmqvist E, Berggren S, Rizvanovic A. 2020. RNA-binding activity and regulatory functions of the emerging sRNA-binding protein ProQ. Biochim Biophys Acta Gene Regul Mech 1863:194596. https://doi.org/10.1016/j.bbagrm.2020.194596.

Jamet A, Jousset AB, Euphrasie D, Mukorako P, Boucharlat A, Ducousso A, Charbit A, Nassif X. 2015. A new family of secreted toxins in pathogenic Neisseria species. PLoS Pathog 11:e1004592. https://doi.org/10.1371/journal.ppat.1004592.

Lalaouna D, Eyraud A, Devinck A, Prévost K, Massé E. 2019. GcvB small RNA uses two distinct seed regions to regulate an extensive targetome. Mol Microbiol 111:473–486. https://doi.org/10.1111/mmi.14168.

Langmead B, Salzberg SL. 2012. Fast gapped-read alignment with Bowtie 2. Nat Methods 9:357–359. https://doi.org/10.1038/nmeth.1923.

Liao Y, Smyth GK, Shi W. 2014. featureCounts: an efficient general purpose program for assigning sequence reads to genomic features. 30:923–930. https://doi.org/10.1093/bioinformatics/btt656.

Livak KJ, Schmittgen TD. 2001. Analysis of relative gene expression data using real-time quantitative PCR and the 2(-Delta Delta C(T)) method. Methods 25:402–408. https://doi.org/10.1006/meth.2001.1262.

Love MI, Huber W, Anders S. 2014. Moderated estimation of fold change and dispersion for RNA-seq data with DESeq2. Genome Biol 15:550. https://doi.org/10.1186/s13059-014-0550-8.

Mann M, Wright PR, Backofen R. 2017. IntaRNA 2.0: enhanced and customizable prediction of RNA-RNA interactions. Nucleic Acids Res 45:W435–W439. https://doi.org/10.1093/nar/gkx279.

Martin M. 2011. Cutadapt removes adapter sequences from high-throughput sequencing reads. EMBnet.journal 17:10–12. https://doi.org/10.14806/ej.17.1.200.

Mehlitz A, Eylert E, Huber C, Lindner B, Vollmuth N, Karunakaran K, Goebel W, Eisenreich W, Rudel T. 2017. Metabolic adaptation of Chlamydia trachomatis to mammalian host cells. Mol Microbiol 103:1004–1019. https://doi.org/10.1111/mmi.13603.

Miller HK, Auerbuch V. 2015. Bacterial iron-sulfur cluster sensors in mammalian pathogens. Metallomics 7:943–956. https://doi.org/10.1039/c5mt00012b.

Miyakoshi M, Chao Y, Vogel J. 2015. Cross talk between ABC transporter mRNAs via a target mRNA-derived sponge of the GcvB small RNA. EMBO J 34:1478–1492. https://doi.org/10.15252/embj.201490546.

Miyakoshi M, Okayama H, Lejars M, Kanda T, Tanaka Y, Itaya K, Okuno M, Itoh T, Iwai N, Wachi M. 2021. Mining RNA-seq data reveals the massive regulon of GcvB small RNA and its physiological significance in maintaining amino acid homeostasis in Escherichia coli. Mol Microbiol 117:160–178. https://doi.org/10.1111/mmi.14814.

Mollerup MS, Ross JA, Helfer AC, Meistrup K, Romby P, Kallipolitis BH. (2016). Two novel members of the LhrC family of small RNAs in Listeria monocytogenes with overlapping regulatory functions but distinctive expression profiles. RNA Biol 13:895–915. https://doi.org/10.1080/15476286.2016.1208332.

Monaco C, Talà A, Spinoas MR, Progida C, De Nitto E, Gaballo A, Bruni CB, Bucci C, Alifano P. 2006. Identification of a meningococcal L-glutamate ABC transporter operon essential for growth in low-sodium environments. Infect Immun 74:1725–1740. https://doi.org/10.1128/IAI.74.3.1725-1740.2006.

Morgenstern J, Campos Campos M, Nawroth P, Fleming T. 2020. The glyoxalase system-new insights into an ancient metabolism. Antioxidants (Basel) 9:939. https://doi.org/10.3390/antiox9100939.

Morse SM, Stein S, Hines J. 1974. Glucose metabolism in Neisseria gonorrhoeae. J Bacteriol 120:702–714. https://doi.org/10.1128/jb.120.2.702-714.1974.

Overton TW, Justino MC, Li Y, Baptista JM, Melo AM, Cole JA, Saraiva LM. 2008. Widespread distribution in pathogenic bacteria of di-iron proteins that repair oxidative and nitrosative damage to iron-sulfur centers. J Bacteriol 190:2004–2013. https://doi.org/10.1128/JB.01733-07.

Overton TW, Whitehead R, Li Y, Snyder LA, Saunders NJ, Smith H, Cole JA. 2006. Coordinated regulation of the Neisseria gonorrhoeae-truncated denitrification pathway by the nitric oxide-sensitive repressor, NsrR, and nitrite-insensitive NarQ-NarP. J Biol Chem 281:33115–33126. https://doi.org/10.1074/jbc.M607056200.

Pannekoek Y, Huis In ‘t Veld RA, Schipper K, Bovenkerk S, Kramer G, Brouwer MC, van de Beek D, Speijer D, van der Ende A. 2017. Neisseria meningitidis uses sibling small regulatory RNAs to switch from cataplerotic to anaplerotic metabolism. mBio 8:e02293–16. https://doi.org/10.1128/mBio.02293-16.

Papenfort K, Vanderpool CK. 2015. Target activation by regulatory RNAs in bacteria. FEMS Microbiol Rev 39:362–378. https://doi.org/10.1093/femsre/fuv016.

Potts AH, Vakulskas CA, Pannuri A, Yakhnin H, Babitzke P, Romeo T. 2017. Global role of the bacterial post-transcriptional regulator CsrA revealed by integrated transcriptomics. Nat Commun 8:1596. https://doi.org/10.1038/s41467-017-01613-1.

Pulvermacher SC, Stauffer LT, Stauffer GV. 2009. Role of the Escherichia coli Hfq protein in GcvB regulation of oppA and dppA mRNAs. Microbiology (Reading) 155:115–123. https://doi.org/10.1099/mic.0.023432-0.

Quereda JJ, Ortega AD, Pucciarelli MG, García-Del Portillo F. 2014. The Listeria small RNA Rli27 regulates a cell wall protein inside eukaryotic cells by targeting a long 5’-UTR variant. PLoS Genet 10:e1004765. https://doi.org/10.1371/journal.pgen.1004765.

Ramsey ME, Hackett KT, Kotha C, Dillard JP. 2012. New complementation constructs for inducible and constitutive gene expression in Neisseria gonorrhoeae and Neisseria meningitidis. Appl Environ Microbiol 78:3068–3078. https://doi.org/10.1128/AEM.07871-11.

Ravnikar PD, Somerville RL. 1987. Genetic characterization of a highly efficient alternate pathway of serine biosynthesis in Escherichia coli. J Bacteriol 169:2611–2617. https://doi.org/10.1128/jb.169.6.2611-2617.1987.

Remmele CW, Xian Y, Albrecht M, Faulstich M, Fraunholz M, Heinrichs E, Dittrich MT, Müller T, Reinhardt R, Rudel T. 2014. Transcriptional landscape and essential genes of Neisseria gonorrhoeae. Nucleic Acids Res 42:10579–10595. https://doi.org/10.1093/nar/gku762.

Ribeiro FJ, Przybylski D, Yin S, Sharpe T, Gnerre S, Abouelleil A, Berlin AM, Montmayeur A, Shea TP, Walker BJ, Young SK, Russ C, Nusbaum C, MacCallum I, Jaffe DB. 2012. Finished bacterial genomes from shotgun sequence data. Genome Res 22:2270–2277. https://doi.org/10.1101/gr.141515.112

Rice PA, Shafer WM, Ram S, Jerse AE. 2017. Neisseria gonorrhoeae: drug resistance, mouse models, and vaccine development. Annu Rev Microbiol 71:665–686. https://doi.org/10.1146/annurev-micro-090816-093530.

Sedlyarova N, Shamovsky I, Bharati BK, Epshtein V, Chen J, Gottesman S, Schroeder R, Nudler E. 2016. sRNA-mediated control of transcription termination in E. coli. Cell 167:111–121.e13. https://doi.org/10.1016/j.cell.2016.09.004.

Sharma CM, Papenfort K, Pernitzsch SR, Mollenkopf HJ, Hinton JC, Vogel J. 2011. Pervasive post-transcriptional control of genes involved in amino acid metabolism by the Hfq-dependent GcvB small RNA. Mol Microbiol 81:1144–1165. https://doi.org/10.1111/j.1365-2958.2011.07751.x.

Sharma CM, Darfeuille F, Plantinga TH, Vogel J. 2007. A small RNA regulates multiple ABC transporter mRNAs by targeting C/A-rich elements inside and upstream of ribosome-binding sites. Genes Dev 21:2804–2817. https://doi.org/10.1101/gad.447207.

Sheehan LM, Caswell CC. 2018. An account of evolutionary specialization: the AbcR small RNAs in the Rhizobiales. Mol Microbiol 107:24–33. https://doi.org/10.1111/mmi.13869.

Sievers S, Sternkopf Lillebaek EM, Jacobsen K, Lund A, Mollerup MS, Nielsen PK, Kallipolitis BH. 2014. A multi-copy sRNA of Listeria monocytogenes regulates expression of the virulence adhesin LapB. Nucleic Acids Res 42:9383–9398. https://doi.org/10.1093/nar/gku630.

Sittka A, Pfeiffer V, Tedin K, Vogel J. 2007. The RNA chaperone Hfq is essential for the virulence of Salmonella typhimurium. Mol Microbiol 63:193–217. https://doi.org/10.1111/j.1365-2958.2006.05489.x.

Stauffer GV. 2004. Regulation of serine, glycine, and one-carbon biosynthesis. EcoSal Plus 1. https://doi.org/10.1128/ecosalplus.3.6.1.2.

Steiner TM, Lettl C, Schindele F, Goebel W, Haas R, Fischer W, Eisenreich W. 2021. Substrate usage determines carbon flux via the citrate cycle in Helicobacter pylori. Mol Microbiol 116:841–860. https://doi.org/10.1111/mmi.14775.

Storz G, Vogel J, Wassarman KM. 2011. Regulation by small RNAs in bacteria: expanding frontiers. Mol Cell 43:880–891. https://doi.org/10.1016/j.molcel.2011.08.022.

Thomas MT, Shepherd M, Poole RK, van Vliet AHM, Kelly DJ, Pearson BM. 2011. Two respiratory enzyme systems in Campylobacter jejuni NCTC 11168 contribute to growth on L-lactate. Environ Microbiol 13:48–61. https://doi.org/10.1111/j.1462-2920.2010.02307.x.

Torres-Quesada O, Millán V, Nisa-Martínez R, Bardou F, Crespi M, Toro N, Jiménez-Zurdo JI. 2013. Independent activity of the homologous small regulatory RNAs AbcR1 and AbcR2 in the legume symbiont Sinorhizobium meliloti. PLoS One 8:e68147. https://doi.org/10.1371/journal.pone.0068147.

Torres-Quesada O, Reinkensmeier J, Schlüter JP, Robledo M, Peregrina A, Giegerich R, Toro N, Becker A, Jiménez-Zurdo JI. 2014. Genome-wide profiling of Hfq-binding RNAs uncovers extensive post-transcriptional rewiring of major stress response and symbiotic regulons in Sinorhizobium meliloti. RNA Biol 11:563–579. https://doi.org/10.4161/rna.28239.

Unemo M, Shafer WM. 2014. Antimicrobial resistance in Neisseria gonorrhoeae in the 21st century: past, evolution, and future. Clin Microbiol Rev 27:587–613. https://doi.org/10.1128/CMR.00010-14.

Urbanowski ML, Stauffer LT, Stauffer GV. 2000. The gcvB gene encodes a small untranslated RNA involved in expression of the dipeptide and oligopeptide transport systems in Escherichia coli. Mol Microbiol 37:856–868. https://doi.org/10.1046/j.1365-2958.2000.02051.x.

Vogel J, Luisi BF. 2011. Hfq and its constellation of RNA. Nat Rev Microbiol 9:578–589. https://doi.org/10.1038/nrmicro2615.

Yang Q, Figueroa-Bossi N, Bossi L. 2014. Translation enhancing ACA motifs and their silencing by a bacterial small regulatory RNA. PLoS Genet 10:e1004026. https://doi.org/10.1371/journal.pgen.1004026.

Zachary M, Bauer S, Klepsch M, Wagler K, Hüttel B, Rudel T, Beier D. 2021. Identification and initial characterization of a new pair of sibling sRNAs of Neisseria gonorrhoeae involved in type IV pilus biogenesis. Microbiology (Reading) 167:001080. https://doi.org/10.1099/mic.0.001080.

