## Supplementary figures and images for "A central role of sibling sRNAs NgncR_162/163 in main metabolic pathways of *Neisseria gonorrhoeae*"

### Supplemental Figure 1

#### Slide 1
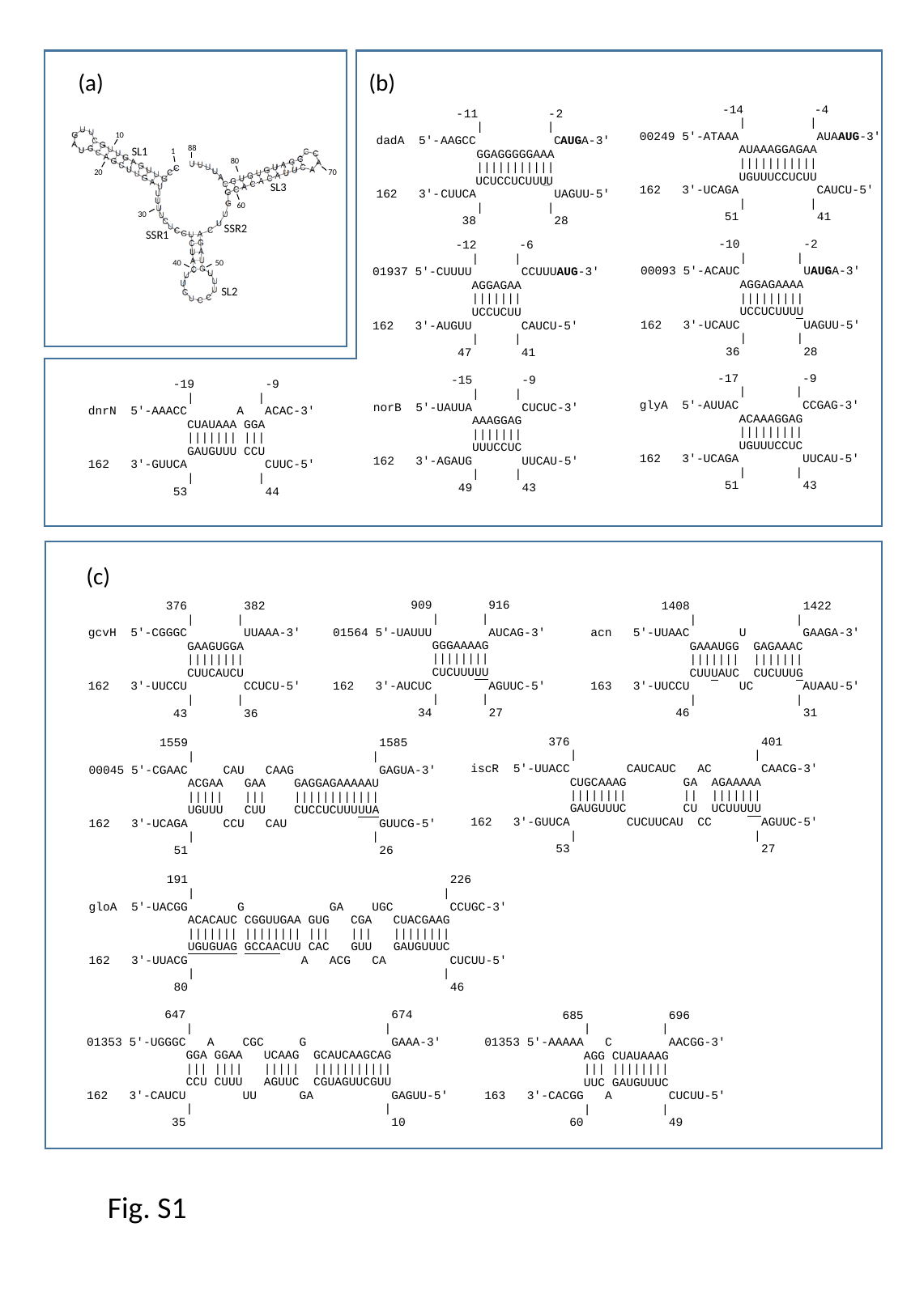

(a)
(b)
-
11
-
2
| |
AUG
dadA
5'
-
AAGCC C
A
-
3'
GGAGGGGGAAA
|||||||||||
UCUCCUCUUU
U
162
3'
-
CUUCA UAGUU
-
5'
| |
38 28
10
88
SL1
1
80
20
70
SL3
60
30
SSR2
SSR1
40
50
SL2
(c)
Fig. S1

### Supplemental Figure 2

#### Slide 1
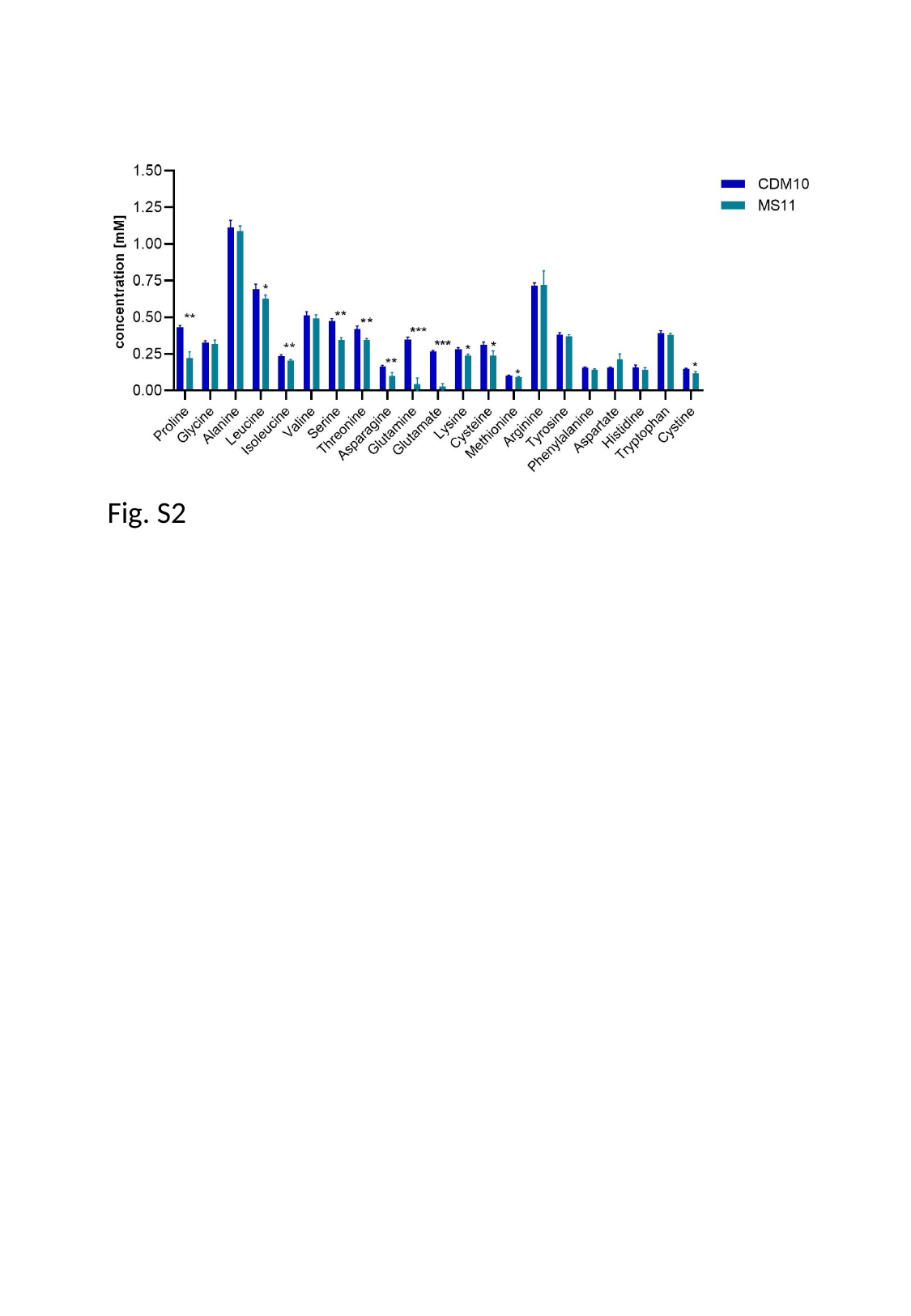

Fig. S2

### Supplemental Figure 3

#### Slide 1
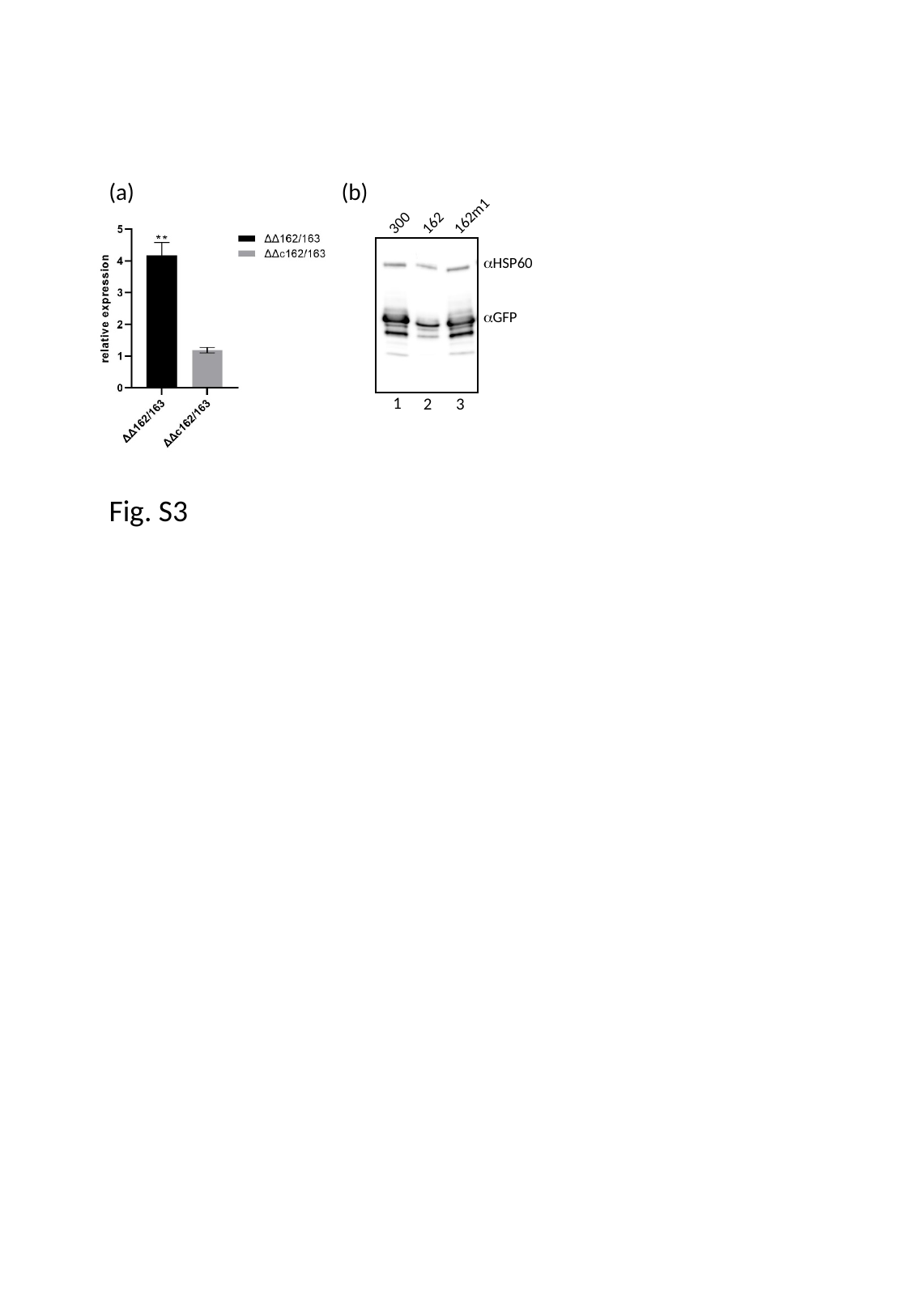

(b)
(a)
162m1
300
162
αHSP60
αGFP
1
2
3
Fig. S3

### Supplemental Figure 4

#### Slide 1
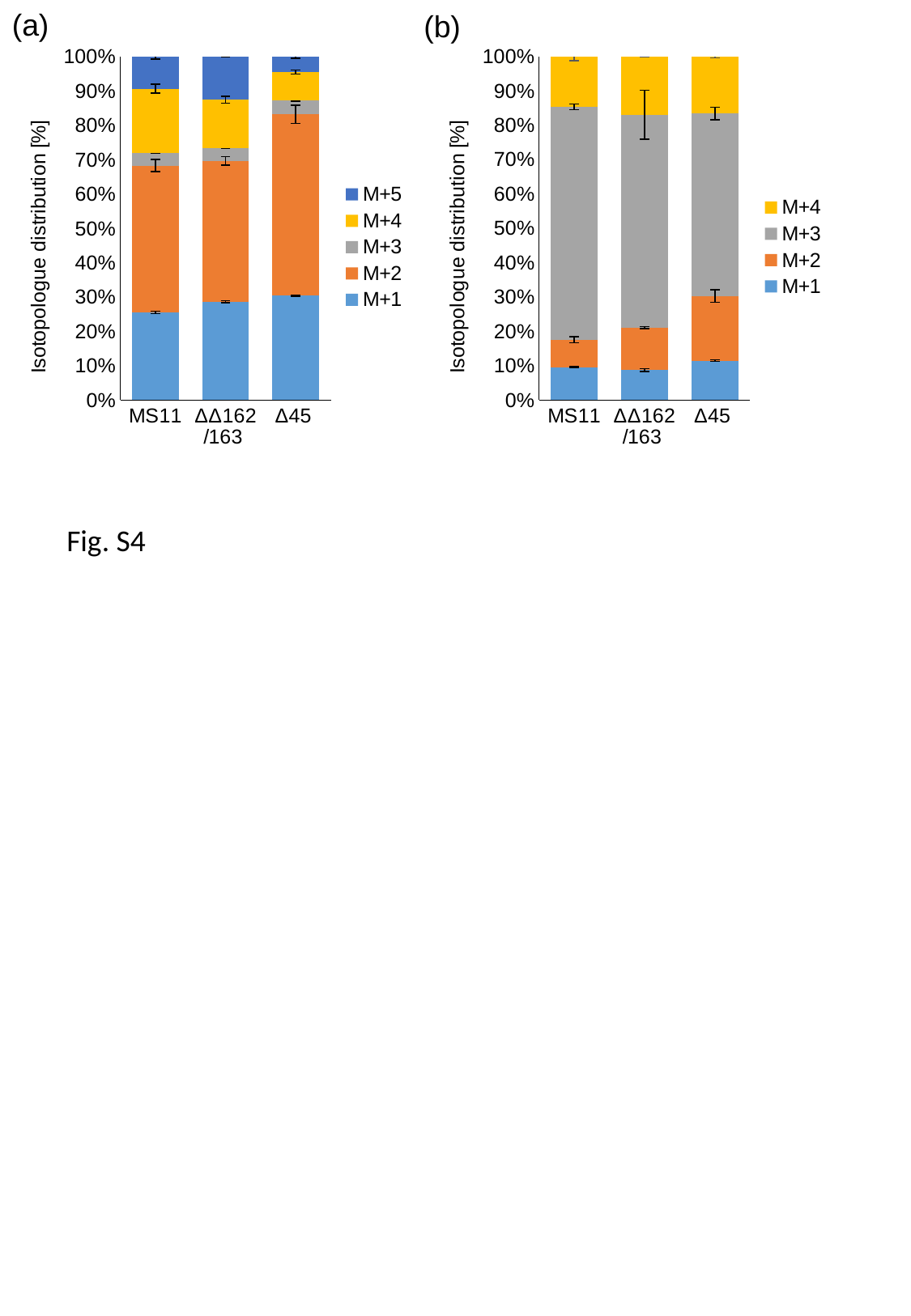

(a)
(b)
##### Chart
| Category | M+1 | M+2 | M+3 | M+4 |
|---|---|---|---|---|
| MS11 | 0.0172358104654846 | 0.014510629005380733 | 0.12206171575091496 | 0.02625214164744405 |
| ΔΔ162/163 | 0.019710179344641768 | 0.027612394279361713 | 0.13890040021570602 | 0.03802022988962298 |
| Δ45 | 0.020212483996492933 | 0.03301565895272344 | 0.09314392565413611 | 0.029094731038026476 |
##### Chart
| Category | M+1 | M+2 | M+3 | M+4 | M+5 |
|---|---|---|---|---|---|
| MS11 | 0.02069434936276283 | 0.03456288357672291 | 0.0029756010706115184 | 0.015123530022057229 | 0.00747812658355125 |
| ΔΔ162/163 | 0.023581128218382263 | 0.03379276957054854 | 0.003065666827208651 | 0.011596937389727453 | 0.010247392779706755 |
| Δ45 | 0.023084610211807063 | 0.04006197733761932 | 0.0030572026110390685 | 0.00625306035219606 | 0.0033161091033410066 |Fig. S4

### Supplemental Figure 5

#### Slide 1
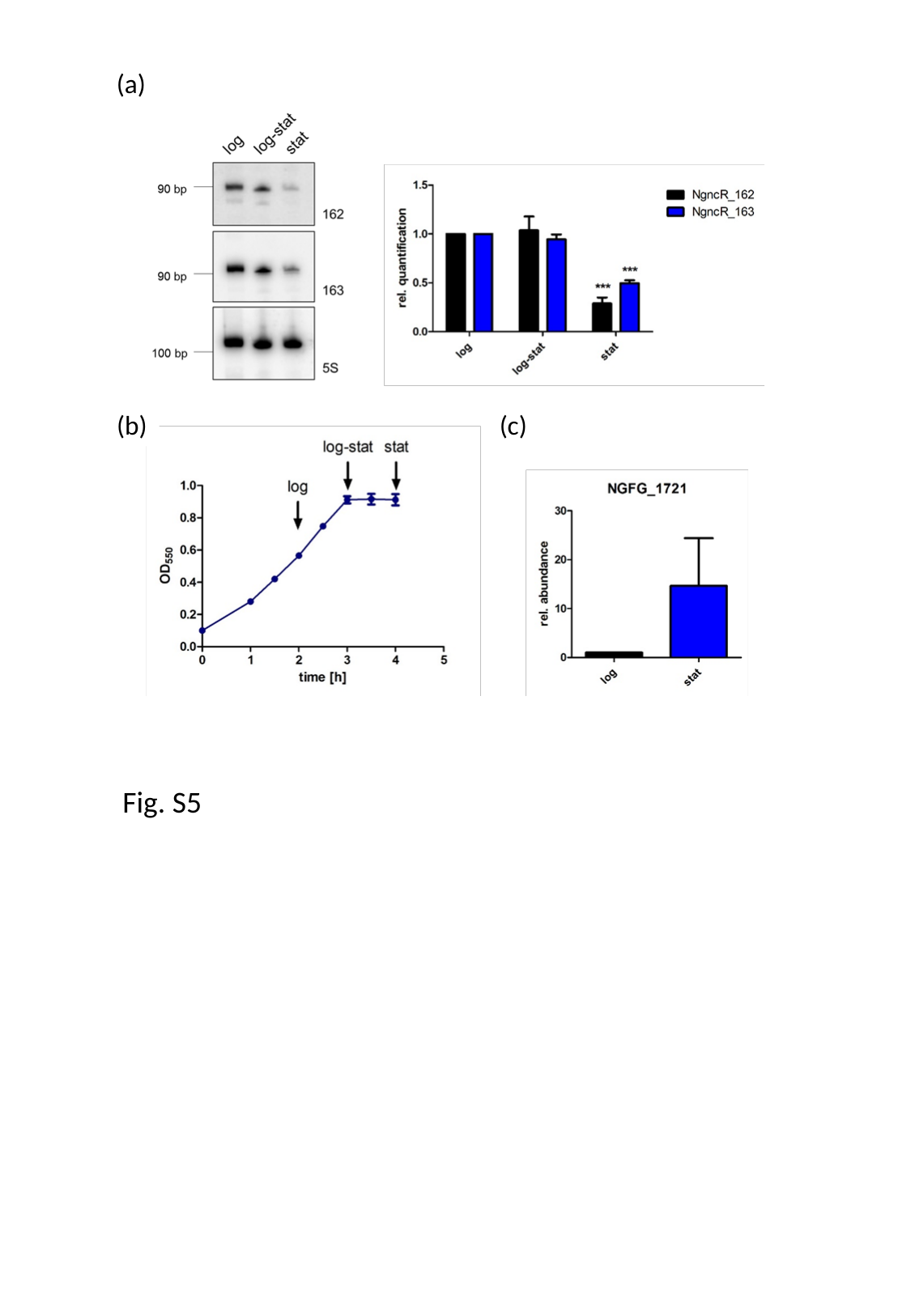

(a)
(b)
(c)
Fig. S5
