## Supplemental Table 1 for "A central role of sibling sRNAs NgncR_162/163 in main metabolic pathways of *Neisseria gonorrhoeae*"

**Table S1: Bacterial strains and plasmids used in this study**

| Strain / plasmid | description | reference |
| --- | --- | --- |
| *E. coli* |  |  |
| DH5α | F- gyrA96 (Nalr) recA1 relA1 endA1 thi-1 hsdR17(rk-mk+) glnV44 deoRΔ(lacZYA-argF)U169 [Φ80dΔ(lacZ)M15] | Thermo Fisher Scientific |
| Top10 | F- mcrA Δ( mrr-hsdRMS-mcrBC) Φ80lacZΔM15 Δ lacX74 recA1 araD139 Δ(araleu)7697 galU galK rpsL (StrR) endA1 nupG | Thermo Fisher Scientific |
| *N. gonorrhoeae* |  |  |
| MS11 | wild-type *N. gonorrhoeae* | laboratory strain  collection |
| MS11 ΔΔ162/163 | MS11 with sRNA genes NgncR_162 and NgncR_163 substituted by a kanamycin resistance cassette | Bauer et al., 2017 |
| MS11 ΔΔc162 | MS11 ΔΔ162/163 with sRNA gene NgncR_162 inserted in *iga-trpB* locus | Bauer et al., 2017 |
| MS11 ΔΔc163 | MS11 ΔΔ162/163 with sRNA gene NgncR_162 inserted in *iga-trpB* locus | Bauer et al., 2017 |
| MS11 ΔΔc162/163 | MS11 ΔΔ162/163 with sRNA genes NgncR_162 and NgncR_163 inserted in *iga-trpB* locus | Bauer et al., 2017 |
| MS11 P*_opa_*45 | MS11 expressing NGFG_00045 under control of the P*_opa_* promoter | this study |
| MS11 P*_opa_*45 ΔΔ162/163 | MS11 P*_opa_*45 with sRNA genes NgncR_162 and NgncR_163 substituted by a kanamycin resistance cassette | this study |
| MS11 P*_opa_*45 ΔΔc162/163 | MS11 P*_opa_*45 ΔΔ162/163 with sRNA genes NgncR_162 and NgncR_163 inserted in *iga-trpB* locus | this study |
| MS11 P_45_gfp | MS11 with *gfp* replacing NGFG_00045 | this study |
| MS11 P_45_gfpSD | MS11 P_45_gfp with an artificial RBS introduced into the NGFG_00045 5’-UTR | this study |
| MS11 ΔΔP_45_gfp | MS11 ΔΔ162/163 with *gfp* replacing NGFG_00045 | this study |
| MS11 ΔΔP_45_gfpSD | MS11 ΔΔP_45_gfp with an artificial RBS introduced into the NGFG_00045 5’-UTR | this study |
| MS11 Δ45 | MS11 with the DNA segment encompassing 289 nucleotides from the upstream region and 239 bp from the 5’-end of NGFG_00045 replaced by *ermC* | this study |
| MS11 Δc45 | derivative of MS11 Δ45 with reintegration of the DNA segment encompassing 289 nucleotides from the upstream region and 239 bp from the 5’-end of NGFG_00045 | this study |
| MS11 Δ1564 | MS11 with NGFG_01564 substituted by *ermC* | this study |
| MS11 Δ1721 | MS11 with NGFG_01721 substituted by *aadA1* | this study |
| plasmids |  |  |
| pSL1180 | cloning vector | Brosius, 1989 |
| pMR68 | complementation plasmid for *N. gonorrhoeae* and *N. meningitidis* | Ramsey et al., 2012 |
| pJV300 | plasmid expressing a nonsense sRNA under control of the P_LlacO_ promoter | Sittka et al., 2007 |
| pXG30-SF | superfolder *gfp*-based translational fusion plasmid for intercistronic fusions | Corcoran et al., 2012 |
| pJV-162 | derivative of pJV300 expressing NgncR_162 | Bauer et al., 2017 |
| pJV-162m1 | derivative of pJV300 expressing NgncR_162 with a mutated SL2 sequence | Bauer et al., 2017 |
| pXG-863 | pXG30-SF derivative expressing a translational *glyA*-*gfp* fusion | this study |

**Tab. S2:** **Oligonuncleotides used in this study**

| Name | Sequence (5‘ to 3‘)^a^ | amplification of |
| --- | --- | --- |
| 45-5UTR-1 | taatgaattcgccgtgctgaaCTCCGCTGCGCC  AAATCGTTGCC | 3’-end of NGFG_00044 and part of intergenic region between NGFG_00044 and NGFG_00045 |
| 45-5UTR-23 | CAATTAACCCTCACTAAAggtaccGG  AAATAACCGAAACCGGACG | 3’-end of NGFG_00044 and part of intergenic region between NGFG_00044 and NGFG_00045 with 3’-overhang to *ermC* |
| 45-5ermC-13 | CGTCCGGTTTCGGTTATTTCCggtacc  TTTAGTGAGGGTTAATTG | *ermC* with 5‘-overhang to upstream region of NGFG_00045 |
| 45-5ermC-23 | CCTTAACAGGGAAAGCAGCAGctgcag  GTACACGAAAAACAAGTT | *ermC* with 3‘-overhang to downstream region of NGFG_00045 |
| ermPopa1 | TGGCGGATTAACAAAAACCGGctgcag  GTACACGAAAAACAAGTT | *ermC* with 3‘-overhang to *opa* promoter (P*_opa_*) |
| ermPopa2 | CTTGTTTTTCGTGTACCTGCAG  CCGGTTTTTGTTAATCCGCCA | P*_opa_* with 5‘-overhang to *ermC* |
| 45Popa-4 | CAATCTATGTGCTTATCGTAAAAA  ATTATATCGGGTTCCGGGCG | P*_opa_* with 3‘-overhang to the 5’-UTR of NGFG_00045 |
| 45Popa-5 | CGCCCGGAACCCGATATAAT  TTTTTACGATAAGCACATAGATTG | 5’-UTR and part of coding region of NGFG_00045 with 5’-overhang to P*_opa_* |
| 45Flag-6 | attatagagctcGTAAAAACCCACACGCCCGCCAA | 5’-UTR and part of coding region of NGFG_00045 |
| 45gfp-1 | tataatgtcgacGCGAATAAGTGCGGCTAAGG | promoter region and 5‘-UTR of NGFG_00045 |
| 45gfp-3 | GTGAAAAGTTCTTCTCCTTTACTCAT ATATAGAAACAGCGTCTGACAGG | promoter region and 5‘-UTR of NGFG_00045 with 5‘-overhang to *gfp*-mut2 |
| 45gfp-2 | CCTGTCAGACGCTGTTTCTATATATGAGT AAAGGAGAAGAACTTTTC | *gfp*-mut2 with 5‘-overhang to 5‘-UTR of NGFG_00045 |
| 45gfp-7 | CAATTAACCCTCACTAAAggtacctctagaGCC  GTCTGAAAACAGCC | *gfp*-mut2 with 3‘-overhang to *ermC* |
| 45gfp-6 | GCTGTTTTCAGACGGCtctagaggtaccTTTAG  TGAGGGTTAATTG | downstream region of NGFG_00045 with 5‘-overhang to ermC |
| 45mut-5 | ATTATAGAGCTCGGGGTGCAATATCTAAGG  AATT | downstream region of NGFG_00045 |
| 45gfp-8 | CCTTTACTCATATGTATATCTCCTTCTGAC AGGGATAATGTCTTC | promoter region and mutated 5‘-UTR of NGFG_00045 with 5‘-overhang to *gfp*-mut2 |
| 45gfp-9 | GAAGACATTATCCCTGTCAGAAGGAGA TATACATATGAGTAAA | *gfp*-mut2 with 5‘-overhang to mutated 5‘-UTR of NGFG_00045 |
| 45-3ermC-3 | CGATGGCATAATCGAGCAGCCTGCAGG TACACGAAAAACAAGTT | *ermC* with 3‘-overhang to coding region of NGFG_00045 |
| D45-1 | CTTGTTTTTCGTGTACCTGCAGGCTGCT  CGATTATGCCATCG | coding region of NGFG_00045 with 5’-overhang to *ermC* |
| D45-2 | ACCATAATGCCGAAGCAGATGG | coding region of NGFG_00045 |
| D45-3 | TGAGACACAATTCATCGATGATGGAAAT AACCGAAACCGGACG | 3’-end of NGFG_00044 and part of intergenic region between NGFG_00044 and NGFG_00045 with 3’-overhang to kan^r^ |
| D45-4 | CGTCCGGTTTCGGTTATTTCCATCATCG ATGAATTGTGTCTCAA | kan^r^ with 5‘-overhang to intergenic region between NGFG_00044 and NGFG_00045 |
| D45-5 | CCAACGCGGCAGGAATCTATCCTGAAG CTTGCATGCCTGCA | kan^r^ with 3‘-overhang to upstrem region of NGFG_00045 |
| D45-6 | TGCAGGCATGCAAGCTTCAGGATAGAT  TCCTGCCGCGTTGG | upstream region of NGFG_00045 with 5‘-overhang to kan^r^ |
| D1564-1 | gccgtctgaaATTGAAGCCTGTGTGATACTGC | upstream region of NGFG_01564 |
| D1564-2 | CGCAATTAACCCTCACTAAAGAAGGCTT TCAGACGGCATAGG | upstream region of NGFG_01564 with 3’-overhang to *ermC* |
| D1564-3 | CCTATGCCGTCTGAAAGCCTTCTTTAGT GAGGGTTAATTGCG | *ermC* with 5’-overhang to upstream region of NGFG_01564 |
| D1564-4 | GGACAGCCAGCGCAATAAAGACGAAAA ACAAGTTAAGGGATGC | *ermC* with 3’-overhang to coding region of NGFG_01564 |
| D1564-5 | GCATCCCTTAACTTGTTTTTCGTCTTTATT GCGCTGGCTGTCC | part of the coding region of NGFG_01564 with 5’-overhang to *ermC* |
| D1564-6 | AAAATAGCCGTGCCGATAAGC | part of the coding region of NGFG_01564 |
| 1721up1 | tataatgagctcTTCCAATTCTGCCGCATCG | upstream region of NGFG_01721 |
| 1721up2 | tataatctgcagAATGCTGCCGACTGCCATACT | upstream region of NGFG_01721 |
| Spec2(PstI) | tataatctgcagTGTAGGGCTTATTATGCAGC | spectionomycin resistance cassette (spec^r^) |
| Popa5(KpnI) | tataatggtaccGGTTTTTGTTAATCCGCCA | spectionomycin resistance cassette (spec^r^) |
| D1721-4 | CGTGCCGTCTGAATATTCGggtaccGGTT  TTTGTTAATCCG | spectinomycin resistance cassette with 3’-overhang to downstream region of NGFG_01721 |
| D1721-3 | CGGATTAACAAAAACCggtaccCGAATA  TTCAGACGGCACGGC | downstream region of NGFG_01721 with 5’-overhang to spec^r^ |
| D1721-2 | tataatggatccGATGACCGTTACTTCATGTCC | downstream region of NGFG_01721 |
| 5UTR863-1 | tataatatgcat TTACGCATTATGGGCTATCTGC | intergenic region and regions encoding the last 8 amino acids (aa)of NGFG_00864 and first 40 aa of NGFG_00863 (*glyA*) |
| 5UTR863-2 | tataatgctagcGCTGACGTAGTTTTCAGAGGC | intergenic region and regions encoding the last 8 amino acids (aa)of NGFG_00864 and first 40 aa of NGFG_00863 (*glyA*) |
| 5S FW | CGGCCATAGCGAGTTGGT | amplicon for 5S RNA transcript quantification |
| 5S RV | TTGGCAGTGACCTACTTTCG | amplicon for 5S RNA transcript quantification and Northern Blot probe for 5S RNA |
| qRT45-1 | TCAGGACAAGCTGAACATCG | amplicon for NGFG_00045 transcript quantification |
| qRT45-2 | TTTGTCCATCACGTCCAAAA | amplicon for NGFG_00045 transcript quantification |
| qRT1146-1 | TGGCGCAACCGTTGATCATA | amplicon for NGFG_01146 *(dnrN)* transcript quantification |
| qRT1146-2 | GCAATTTCCGCCGGAAAGGT | amplicon for NGFG_01146 *(dnrN)* transcript quantification |
| qRT1163-1 | CCTCCCGCACAAATCAACAT | amplicon for NGFG_01163 (*icsR*) transcript quantification |
| qRT1163-2 | AATTCTCCCAAAGGTCGTGC | amplicon for NGFG_01163 (*icsR*) transcript quantification |
| qRT1353-1 | CAACGTCAACAGCTTCCTGA | amplicon for NGFG_01353 transcript quantification |
| qRT1353-2 | CGGTAAAGACCTGCATCCAT | amplicon for NGFG_01353 transcript quantification |
| qRT1407-1 | TACCACCTGTAACGGCATGA | amplicon for NGFG_01407 (*acn*) transcript quantification |
| qRT1407-2 | AGGAAAGCCTGTTTCGCATA | amplicon for NGFG_01407 (*acn*) transcript quantification |
| qRT1722-1 | CAATAAAGAGCGCATGGTCA | amplicon for NGFG_01722 (*dadA*) transcript quantification |
| qRT1722-2 | GCTTCGACTTCTTCGGTTTG | amplicon for NGFG_01722 (*dadA*) transcript quantification |
| qRT2111-1 | CAACTGGGATACGGAACGAT | amplicon for NGFG_02111 (*gloA*) transcript quantification |
| qRT2111-2 | GTTGTGCCGTGTTTCATCAG | amplicon for NGFG_02111 (*gloA*) transcript quantification |
| qRT2263-1 | GGCAAAGTCGGCTACAAAAA | amplicon for NGFG_02263 transcript quantification |
| qRT2263-2 | CCGGAAGCCAAAATAAACAA | amplicon for NGFG_02263 transcript quantification |
| qRT249-1 | CGATGATAGGCGGTTTGATT | amplicon for NGFG_00249 transcript quantification |
| qRT249-2 | CGACCGATAAAAACGTCGTC | amplicon for NGFG_00249 transcript quantification |
| qRT93-1 | TGGACATCAACGTCTTCCAA | amplicon for NGFG_00093 transcript quantification |
| qRT93-2 | GATTTCAGTTTGCCCGGATA | amplicon for NGFG_00093 transcript quantification |
| qRT1564-1 | ACCTCTTGGTTTCCCTGCTT | amplicon for NGFG_01564 transcript quantification |
| qRT1564-2 | GTACCGAACGGCATTTTCAT | amplicon for NGFG_01564 transcript quantification |
| qRT1937-1 | AGAAGCCGCCGATATTGATG | amplicon for NGFG_01937 transcript quantification |
| qRT1937-2 | TATTCATCGCATTGGGCAGC | amplicon for NGFG_01937 transcript quantification |
| qRT1471-1 | ATTGGCGATGGTCAACGAAG | amplicon for NGFG_01471 transcript quantification |
| qRT1471-2 | CCATTCTTCTTTGCTGGTCAAA | amplicon for NGFG_01471 transcript quantification |
| qRT1514-1 | CCGTCGGTATTACCCATCAC | amplicon for NGFG_01514 (*gcvH*) transcript quantification |
| qRT1514-2 | TGCGGCTTTTACATACTCAA | amplicon for NGFG_01514 (*gcvH*) transcript quantification |
| qRT2153-1 | GCCCAACATAAAGATGGCGG | amplicon for NGFG_01514 (*norB*) transcript quantification |
| qRT2153-2 | CTGTGGGTGGAAGGCTTCTT | amplicon for NGFG_01514 (*norB*) transcript quantification |
| qRT2042-1 | ATTTTGTCCGAGATGGTTGC | amplicon for NGFG_01514 (*ilvB*) transcript quantification |
| qRT2042-2 | GATAATTTCGCTGCCGTTGT | amplicon for NGFG_01514 (*ilvB*) transcript quantification |
| qRT863-1 | GTGTGATTTTGTGCCGTGAC | amplicon for NGFG_00863 (*glyA*) transcript quantification |
| qRT863-2 | CGCTTCTTTAAACGCTACGG | amplicon for NGFG_00863 (*glyA*) transcript quantification |
| NBS162 | AATCAAGCTGCATCAAGCAACTCAA | Northern blot probe for NgncR_162 |
| NBS163 | TATTAACTGACTACTCGAACCAGCT | Northern blot probe for NgncR_163 |

(a) Sequences introduced for cloning purposes are given in lower case letters. Restriction sites are underlined.
